# Learning RNA secondary structure from chemical probing with MXfold2

**DOI:** 10.64898/2026.11.18.767577

**Authors:** Rintaro Yamauchi, Sho Tsukiyama, Kengo Sato

**Affiliations:** School of Life Science and Technology, Institute of Science Tokyo, Japan; Future Intelligence Visionary Initiative, Institute of Science Tokyo, Japan; AI-Science Nexus Center, Institute of New Industry Incubation, Institute of Science Tokyo, Japan

## Abstract

Training RNA secondary structure predictors is limited by scarce reference annotations, even as chemical probing produces abundant nucleotide-level measurements. These measurements inform pairing states without identifying base-pairing partners. Here we extend MXfold2, which combines neural and thermodynamic folding scores, to learn from probing profiles and predict RNA secondary structures from sequence alone. We introduce three training objectives. One learns from probing-assisted target structures. The other two use implicit maximum-likelihood estimation to pass surrogate gradients from reactivity-ranking and likelihood losses through discrete folding. Simulations, RNAgym measurements and external structural tests show improved chemical agreement and gains in exact base-pair prediction when chemical supervision is combined with structural pretraining and replay. Probing profiles can thus broaden the range of RNAs that inform sequence-only structure prediction, including RNA populations with few complete structure annotations.

---

RNA secondary structure describes which nucleotides pair and provides a basis for understanding RNA folding and function^1^. Consecutive base pairs form stems, and unpaired regions include hairpin and internal loops. Pseudoknots contain base pairs that cross when drawn as arcs above the sequence. Predicting RNA secondary structure from sequence requires choosing a consistent set of partners from many alternatives ^1,2^.

Thermodynamics-based and machine-learning-based methods use different information to score candidate RNA structures. Thermodynamics-based methods sum nearest-neighbor free energy contributions from hairpin loops, internal loops, and base-pair stackings, then use dynamic program- ming to find a minimum-free-energy structure ^2,3^. Machine-learning-based methods learn folding scores from sequences and reference structures. MXfold2 combines a deep neural network’s folding scores with Turner’s nearest-neighbor free energy parameters and uses dynamic programming to select the highest-scoring structure ^1^.

Chemical probing measures how readily a reagent modifies individual nucleotides in folded RNA in vitro or in cells. Selective 2*^′^*-hydroxyl acylation analyzed by primer extension (SHAPE) probes local backbone flexibility at the ribose 2*^′^*-hydroxyl^4,5^. RNAgym includes measurements with the SHAPE reagents 2A3, 1M7, and NMIA^6^. Dimethyl sulfate (DMS) probes base accessibility. Conventional readouts primarily detect modifications on the Watson–Crick pairing faces of adenine (A) and cytosine (C)^7^.

A reactivity profile summarizes the modification signal at each nucleotide. Primer-extension assays locate modifications from stops during reverse transcription. In mutational profiling (MaP), reverse transcription records modifications as sequence changes in complementary DNA. Sequencing and background correction yield nucleotide-level reactivities, as in SHAPE-MaP and DMS-MaPseq ^5,7^.

Reactivity profiles constrain local pairing states but leave pairing partners unresolved. Higher reactivity often indicates flexibility or accessibility. Low reactivity can reflect base pairing, stacking, tertiary contacts, or protection by proteins, and the pattern can depend on cellular conditions^4,8^. Conventional profiles also average measurements across molecules. Even exact knowledge of which positions are paired would permit alternative ways to connect those positions (Figure 1a). A predictor must therefore combine local chemical evidence with other information to select partners.

**Figure 1:**
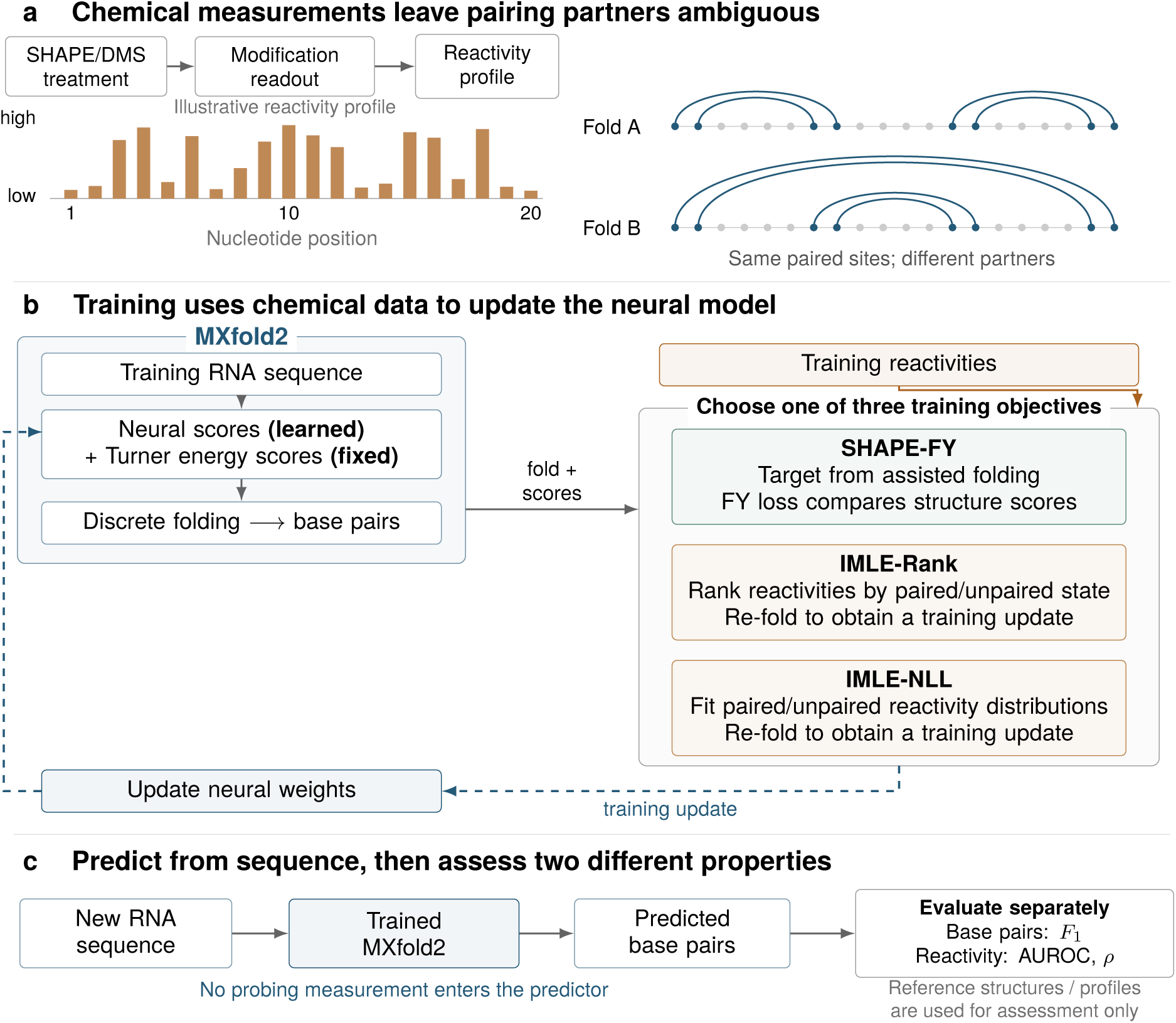
Chemical supervision for sequence-only RNA secondary structure prediction. **a**, Chemical probing with selective 2*^′^*-hydroxyl acylation analyzed by primer extension (SHAPE) or dimethyl sulfate (DMS) produces nucleotide-level reactivity profiles. The profile and folds are schematic: arcs denote base pairs and blue dots denote paired sites. Identical paired sites can have different partners. **b**, SHAPE-FY (SHAPE-guided Fenchel–Young training) learns from targets obtained by assisted folding using a Fenchel–Young loss ^9,17^. Implicit maximum-likelihood estimation (I-MLE) supports direct chemical losses on predicted pairing states^18^: IMLE-Rank (Mann–Whitney ranking) or IMLE-NLL (negative log likelihood). The return arrow denotes learning of neural folding scores. **c**, Prediction after training requires sequence alone. Base-pair *F*_1_ measures partner accuracy; the area under the receiver operating characteristic curve (AUROC) and Spearman correlation measure agreement between predicted unpaired states and reactivity.

*Assisted folding* uses reactivities to guide structure prediction for the RNA that was probed. Measurements can exclude particular pairs or modify their energy contributions. A measurement- dependent energy correction is called a *pseudoenergy* ; for example, high reactivity can make pairing at a position less favorable. Mathews et al.^3^ incorporated chemical-modification constraints into thermodynamics-based methods, and Deigan et al.^9^ introduced logarithmic SHAPE corrections to stacking-related energies. Other approaches fit reactivities to distributions over structures^10,11^. Restrained MaxExpect uses separate paired and unpaired reactivity distributions to derive structural restraints^12^. These approaches require measurements for each RNA being predicted.

Probing profiles can broaden the range of RNAs available for training. High-throughput experi- ments yield nucleotide-level measurements for many RNAs whose pairing partners are unknown, including primary microRNAs and regions of human messenger RNAs ^5–7,13^. Such profiles provide information about which sequence features favor paired or unpaired states. A model trained with these profiles can use that information to predict structures for other RNAs from sequence alone.

Probabilistic folding models show how probing profiles can supervise parameter learning. CONTRAfold-SE learns folding parameters from probing data while treating the unknown structure as a latent variable ^14^. EternaFold extends this approach to joint training on reference structures, chemical mapping, and riboswitch measurements^15^. Its observation model describes reactivities conditional on nucleotide identity and pairing state. It averages the likelihood of the reactivities over structures weighted by the folding model.

Learning from reactivities with MXfold2 requires connecting this incomplete supervision to a discrete structure prediction. The original MXfold2 uses a structured hinge loss that compares a complete reference structure with competing predictions^1^. A reactivity profile provides indirect information about pairing states. The states in the selected structure change discretely with the folding scores, so a loss on those states cannot be backpropagated through structure selection by ordinary differentiation.

Here, we investigate how indirect chemical measurements can train the neural component of MXfold2 while its structures are selected by discrete optimization. We introduce three training objectives for chemical weak supervision. SHAPE-FY uses a Fenchel–Young (FY) loss to learn from probing-assisted structures. IMLE-Rank and IMLE-NLL use implicit maximum-likelihood estimation (I-MLE) to learn from reactivity ranks and distributions. We first generate synthetic profiles from known structures to test this supervision under controlled conditions^16^. We then train with experimental profiles from RNAgym and collections released with eFold, and compare with EternaFold and other released predictors ^6,13,15^. The results show when chemical information can complement complete structure annotations to improve sequence-only prediction.

## Results

### Overview of chemical weak supervision

Our extension trains the neural folding scores of MXfold2 from reactivity profiles and predicts new structures from sequence alone (Figure 1). A profile constrains which positions are paired but cannot identify their partners (Figure 1a). Structural pretraining first supplies partner information from annotated examples; structural replay interleaves such examples with chemical examples during subsequent training.

Each model is trained with one of three alternative chemical objectives (Figure 1b). SHAPE-FY first predicts a target structure using both the current model and the training reactivities, then trains the sequence-only model to favor that target with a Fenchel–Young (FY) loss ^17^. The target is recalculated as the model changes. IMLE-Rank instead encourages predicted unpaired positions to have higher reactivities than paired positions. IMLE-NLL minimizes the negative log likelihood of reactivities under separate paired and unpaired distributions based on the Wu model^12^. The latter two objectives use implicit maximum-likelihood estimation (I-MLE) to pass a surrogate gradient through discrete structure prediction^18^. After training, each model predicts a new RNA from its sequence alone (Figure 1c).

### Learning from simulated probing profiles

The simulations test whether reactivities generated from known structures can improve base-pair prediction after structural pretraining. We pretrain on TrainSetA (3,166 RNAs), generate 2A3 and DMS profiles from TrainSetB reference structures (1,094 RNAs), and evaluate base pairs on TestSetB (430 RNAs) ^1,16^. TrainSetB and TestSetB draw from the same 22 Rfam families (Supplementary Note 2).

All adaptation runs start from the same TrainSetA model pretrained for 100 epochs and add 100 epochs on TrainSetB. IMLE-Rank, IMLE-NLL, and SHAPE-FY learn from simulated TrainSetB reactivity profiles. Two FY controls learn from fixed structures predicted once with RNAfold or MXfold2 using those profiles, and a third learns from the annotated TrainSetB structures. Structural replay interleaves TrainSetA examples with TrainSetB examples. Its percentage is the fraction of all training examples supplied by TrainSetA. The ten method–probe combinations and the reference-structure control each use five replay rates (0, 5, 10, 20, and 50%), giving 55 runs (Figure 2 and Supplementary Table 5).

**Figure 2:**
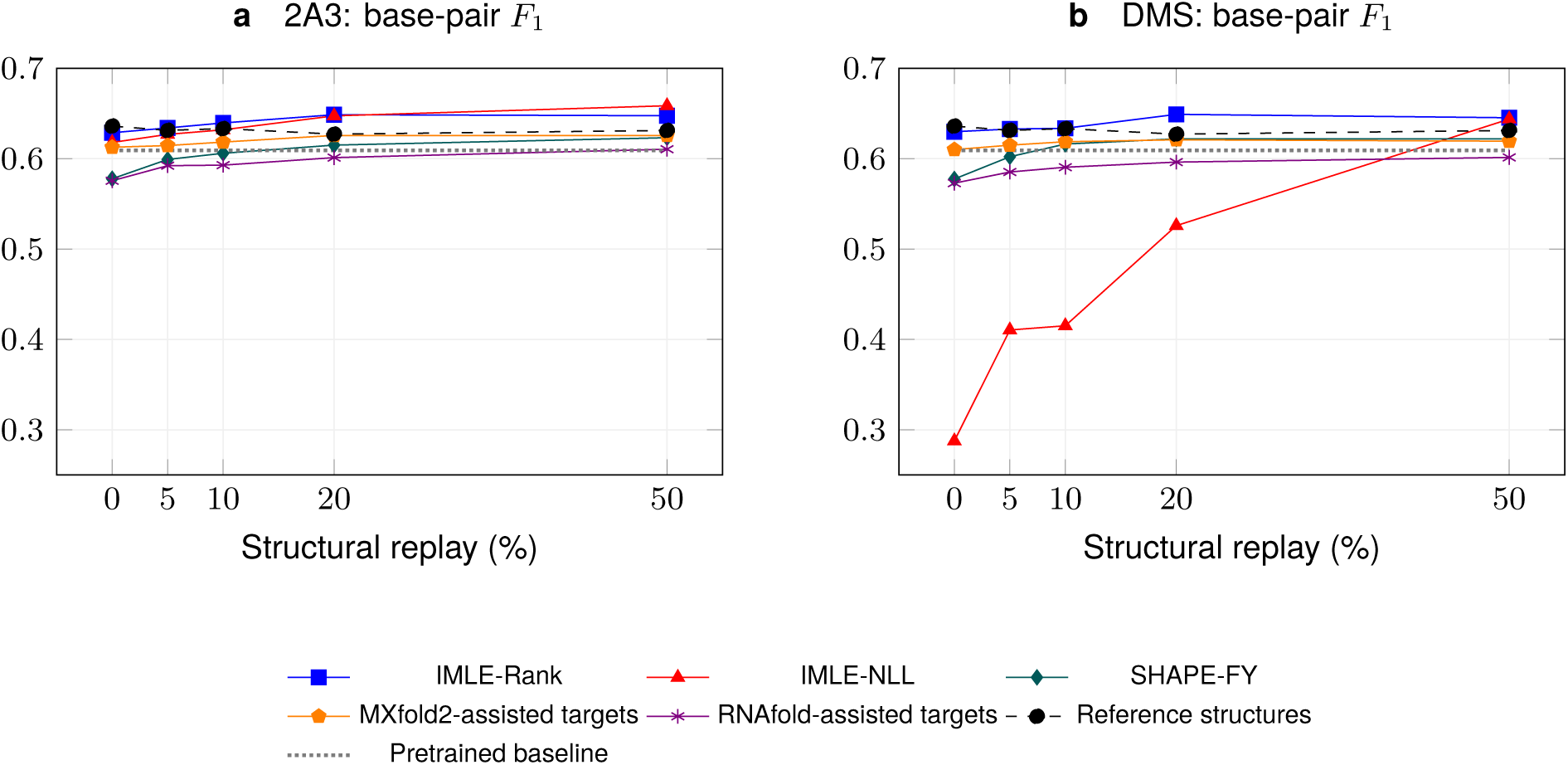
Effect of structural replay on TestSetB accuracy after 100 additional training epochs. **a**, Synthetic profiles for the SHAPE reagent 2A3. **b**, Synthetic dimethyl sulfate (DMS) profiles. Symbols distinguish the five chemical or assisted-target methods and the control trained on reference structures. The horizontal line marks the pretrained baseline. Reactivities are generated from TrainSetB annotations; all runs start from the same TrainSetA model. Larger replay fractions add both structure-labeled examples and optimizer updates. TrainSetB and TestSetB contain related RNAs.

Adding 20% structural replay improved all three chemical objectives relative to their runs without replay. Without replay, IMLE-Rank achieved TestSetB *F*_1_ = 0.629 for 2A3 and 0.630 for DMS, compared with the 0.609 baseline. At 20% replay, both scores increased and rounded to 0.649 at three significant figures. Without replay, the two SHAPE-FY scores fell below the baseline and both rounded to 0.578. At 20% replay, SHAPE-FY reached 0.615 for 2A3 and 0.622 for DMS. IMLE-NLL depended strongly on the probe: 2A3 gave 0.618 without replay and 0.647 at 20%, whereas DMS gave 0.288 and 0.526. DMS IMLE-NLL reached 0.644 only at 50% replay. These differences reflect the full training configuration, including the likelihood, observation filters, clipping, initialization, and optimizer; they do not establish that DMS itself is unsuitable for learning.

IMLE-NLL with synthetic 2A3 and 50% replay achieved the highest final score in this series, *F*_1_ = 0.659. Training directly on the reference structures without replay reached *F*_1_ = 0.636. These endpoints compare complete training configurations, including the loss and replay budget.

Subsequent experiments use 20% structural replay as a common setting, selected using TestSetB results from the TrainSetB SHAPE (2A3) simulations. At that setting, IMLE-Rank improved TestSetB *F*_1_ both for RNAs with a detected TrainSetB match and for those without one. The gain was larger with a match for 2A3 and similar between the two groups for DMS (Supplementary Note 2).

The fixed-teacher controls provided an additional comparison of training targets. At 20% replay, fixed MXfold2 teachers gave *F*_1_ = 0.626 for 2A3 and 0.621 for DMS, while fixed RNAfold teachers gave 0.601 and 0.596, respectively. Increasing replay adds optimizer updates as well as structural supervision.

With synthetic 2A3 profiles, pretraining followed by adaptation gives higher TestSetB accuracy than joint training from random weights for the two I-MLE objectives (Table 1). After 100 joint- training epochs with TrainSetA structures and synthetic TrainSetB 2A3 profiles, TestSetB *F*_1_ reaches 0.613 for IMLE-Rank, 0.635 for IMLE-NLL, and 0.621 for SHAPE-FY. The corresponding pretrained models adapted with 20% structural replay achieve 0.649 and 0.647 for the two I-MLE objectives. SHAPE-FY instead gives a small increase with joint training over its adaptation score of 0.615. The protocols differ in prior structural exposure, observation filters, and optimizer-update budgets. Supplementary Note 2.1 and Table 6 specify these differences and the fixed epoch-100 results for the twelve joint conditions.

**Table 1:** TestSetB accuracy after training with synthetic SHAPE (2A3) profiles. Positive predictive value (PPV) is precision, Sen is sensitivity, and *F*_1_ is averaged over 430 test RNAs. TrainSetA contains 3,166 structure-labeled RNAs and TrainSetB contains 1,094 RNAs with synthetic profiles. The first block adapts a structure-pretrained MXfold2 model for 100 further epochs with 20% structural replay. Fixed assisted-target controls and the reference-structure control use FY. The second block jointly trains MXfold2 on TrainSetA structures and TrainSetB profiles for 100 epochs from random weights. The final block compares EternaFold trained on TrainSetA structures alone with EternaFold jointly trained on those structures and TrainSetB profiles. Both EternaFold models start from zero-valued parameters. Methods and Supplementary Notes 2.1 and 3 give the training and decoding settings.

|  |  | TestSetB |  |  |
|---|---|---|---|---|
| Model | Training supervision | PPV | Sen | *F*<sub>1</sub> |
| *MXfold2: structural pretraining followed by adaptation* |  |  |  |  |
| MXfold2 | Pretrained baseline | 0.579 | 0.660 | 0.609 |
| MXfold2 | IMLE-Rank | 0.630 | 0.687 | 0.649 |
| MXfold2 | IMLE-NLL | 0.642 | 0.670 | 0.647 |
| MXfold2 | SHAPE-FY | 0.579 | 0.672 | 0.615 |
| MXfold2 | RNAfold-assisted targets | 0.566 | 0.658 | 0.601 |
| MXfold2 | MXfold2-assisted targets | 0.593 | 0.679 | 0.626 |
| MXfold2 | Reference structures | 0.616 | 0.655 | 0.627 |
| *MXfold2: joint training from random weights* |  |  |  |  |
| MXfold2 | IMLE-Rank | 0.578 | 0.669 | 0.613 |
| MXfold2 | IMLE-NLL | 0.605 | 0.684 | 0.635 |
| MXfold2 | SHAPE-FY | 0.586 | 0.675 | 0.621 |
| *EternaFold: structural-only control and joint training* |  |  |  |  |
| EternaFold | Structures only | 0.518 | 0.653 | 0.572 |
| EternaFold | Synthetic 2A3 joint | 0.516 | 0.650 | 0.569 |

With EternaFold, joint training on synthetic 2A3 profiles slightly reduced TestSetB accuracy. We fitted the structural-only and joint models separately from zero-valued parameters, using the same 3,166 annotated TrainSetA RNAs in both (Supplementary Note 3). Adding synthetic 2A3 profiles for the 1,094 TrainSetB RNAs yielded a TestSetB *F*_1_ of 0.569, compared with 0.572 for the structural-only model (Table 1).

### Learning from RNAgym profiles

To test learning from experimental profiles, we fine-tuned the TrainSetA model on RNAgym for three epochs with 20% structural replay. We evaluated base-pair accuracy on TestSetB and chemical agreement on the RNAgym test set (Table 2). At the sequence-screen thresholds described in Methods, 5 of the 430 TestSetB RNAs matched the official RNAgym training collection, compared with 125 that matched TrainSetB. From the pretrained TestSetB *F*_1_ of 0.609, IMLE-Rank rose to 0.613, while IMLE-NLL and SHAPE-FY fell to 0.594 and 0.603, respectively.

**Table 2:** Structural accuracy and chemical agreement after RNAgym training. TestSetB columns report mean precision (PPV), sensitivity (Sen), and *F*_1_ over 430 RNAs. AUROC and Spearman correlation *ρ* evaluate predicted unpaired states against 2A3 and DMS test reactivities. The MXfold2 block compares the pretrained model with three models adapted for three epochs with 20% replay, using one profile per RNA and probe. Both EternaFold conditions use all 3,166 TrainSetA RNAs and are evaluated with all retained replicate profiles for these two probes. Methods and Supplementary Table 7 specify their observation sets and aggregation.

|  | TestSetB |  |  | 2A3 |  | DMS |  |
|---|---|---|---|---|---|---|---|
| Training supervision | PPV | Sen | *F*<sub>1</sub> | AUROC | *ρ* | AUROC | *ρ* |
| *MXfold2: one profile per RNA and probe* |  |  |  |  |  |  |  |
| MXfold2 pretrained | 0.579 | 0.660 | 0.609 | 0.696 | 0.332 | 0.762 | 0.418 |
| IMLE-Rank | 0.609 | 0.638 | 0.613 | 0.730 | 0.387 | 0.781 | 0.436 |
| IMLE-NLL | 0.617 | 0.600 | 0.594 | 0.734 | 0.366 | 0.776 | 0.411 |
| SHAPE-FY | 0.575 | 0.650 | 0.603 | 0.712 | 0.357 | 0.770 | 0.419 |
| *EternaFold: all replicate profiles for 2A3 and DMS* |  |  |  |  |  |  |  |
| EternaFold: structures only | 0.518 | 0.653 | 0.572 | 0.689 | 0.317 | 0.754 | 0.399 |
| EternaFold: RNAgym joint | 0.503 | 0.658 | 0.565 | 0.682 | 0.303 | 0.751 | 0.396 |

All three MXfold2 models improved AUROC for both 2A3 and DMS on the RNAgym test set (Table 2 and Supplementary Table 7). The 2A3 AUROC increased from 0.696 to 0.730 for IMLE-Rank, 0.734 for IMLE-NLL, and 0.712 for SHAPE-FY. For IMLE-NLL, DMS Spearman correlation decreased from 0.418 to 0.411.

With EternaFold, joint training on TrainSetA structures and RNAgym profiles reduced mean TestSetB *F*_1_ from 0.572 to 0.565 (Table 2). The joint model also gave lower AUROC and Spearman correlation for 2A3 and DMS on its model-specific observation sets (Supplementary Table 7).

### Fine-tuning after bpRNA and RNAcentral pretraining

To evaluate chemical fine-tuning after broader structural pretraining, we started all three objectives from the same MXfold2 model. We pretrained it for five epochs on bpRNA reference structures^19^ and RNAcentral sequences^20^ labeled with RNAstructure predictions^21^. Fine-tuning then used three epochs on Ribonanza profiles^22^, followed by three epochs on primary microRNA (pri-miRNA) and human messenger RNA (mRNA) profiles (Supplementary Note 4). These profiles come from the collections released with eFold, which uses structures derived from probing-assisted folding as training labels ^13^.

We evaluated the pretrained model and its three fine-tuned models on all 3,795 structural test RNAs released with eFold. The four collections comprise 355 Protein Data Bank (PDB) RNAs ^23^, 3,370 ArchiveII RNAs ^24^, 40 viral fragments^25–28^, and 30 long noncoding RNA (lncRNA) fragments^13,29^. Figure 3 and Supplementary Table 9 report precision, sensitivity, and mean *F*_1_. We screened for sequence overlap to assess whether these tests contain training examples or closely related sequences. The four collections share no full-length exact matches with the pretraining or chemical inputs. Local sequence similarity remains in the PDB and ArchiveII comparisons (Supplementary Note 6).

**Figure 3:**
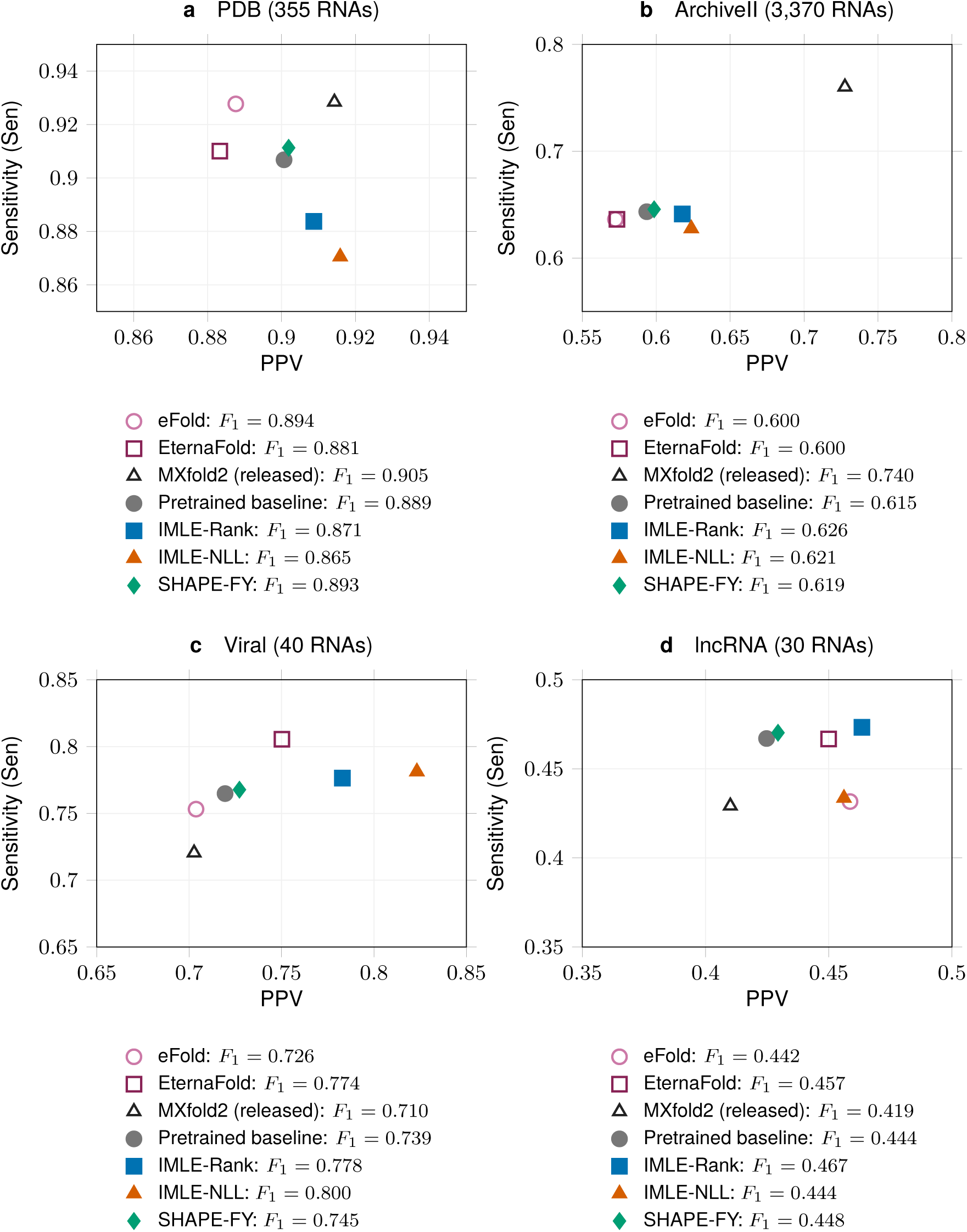
Structural accuracy on the 3,795 test RNAs released with eFold. Points show model-level mean precision (PPV) and sensitivity (Sen); legends give mean per-RNA exact-pair *F*_1_. **a**, Protein Data Bank (PDB). **b**, ArchiveII. **c**, Viral fragments. **d**, Long noncoding RNA (lncRNA). The pretrained baseline uses five bpRNA–RNAcentral epochs. Each chemically trained model adds three Ribonanza epochs and three pri-miRNA/human mRNA epochs. Released eFold, EternaFold, and MXfold2 retain their original training and decoding. Panels use different axis limits.

Averaged over all 3,795 RNAs, mean *F*_1_ increased from 0.640 to 0.649 for IMLE-Rank, 0.645 for IMLE-NLL, and 0.644 for SHAPE-FY. This mean gives equal weight to each RNA. All three objectives improved ArchiveII and viral accuracy from the shared starting model. Mean per-RNA *F*_1_ on ArchiveII increased from 0.615 to 0.626 for IMLE-Rank, 0.621 for IMLE-NLL, and 0.619 for SHAPE-FY. Viral *F*_1_ increased from 0.739 to 0.778, 0.800, and 0.745, respectively. On PDB, IMLE-Rank and IMLE-NLL reduced *F*_1_ from 0.889 to 0.871 and 0.865. Both I-MLE objectives increased PDB precision and reduced sensitivity. IMLE-NLL gave nearly the same lncRNA *F*_1_ as the pretrained model. SHAPE-FY gave small improvements on all four collections.

The released predictors were evaluated on the same test RNAs with their original training histories and decoding settings (Supplementary Note 5.2). PDB *F*_1_ is 0.894 for eFold, 0.881 for EternaFold predictions distributed with the eFold study, and 0.905 for released MXfold2. The released MXfold2 training collection contains full-length exact matches to 97 PDB and 777 ArchiveII references.

### Single-structure chemical agreement

We evaluated chemical agreement on the same 40 viral RNAs used to assess structural prediction (Figure 3c and Supplementary Table 8). IMLE-NLL gave the highest AUROC and Spearman correlation among the four models sharing the five-epoch checkpoint. Its AUROC increased from 0.810 to 0.845 and Spearman correlation from 0.517 to 0.579. IMLE-Rank and SHAPE-FY also increased both overall measures. The combined mean weights all 40 RNAs equally, including 25 measured with DMS and 15 with SHAPE. SHAPE-FY improved chemical agreement on the SHAPE profiles and slightly reduced both DMS measures.

### External structural tests

We tested the same pretrained model and its three fine-tuned models on two additional structural benchmarks, Archive-NoFam and bpRNAnew. Archive-NoFam contains 89 RNAs in 47 structural groups. The primary evaluation excludes seven groups flagged by the benchmark authors for overlap with bpRNA ^30^. On the resulting 79 RNAs in 40 groups (Figure 4a), mean *F*_1_ across groups rose from 0.752 to 0.759 for IMLE-Rank and 0.765 for IMLE-NLL, while SHAPE-FY fell to 0.751. Both I-MLE objectives increased precision and reduced sensitivity on this collection. On all 89 RNAs, IMLE-Rank and IMLE-NLL fell from the pretrained-model score of 0.764 to 0.758 and 0.759, while SHAPE-FY rose to 0.770 (Supplementary Table 11).

**Figure 4:**
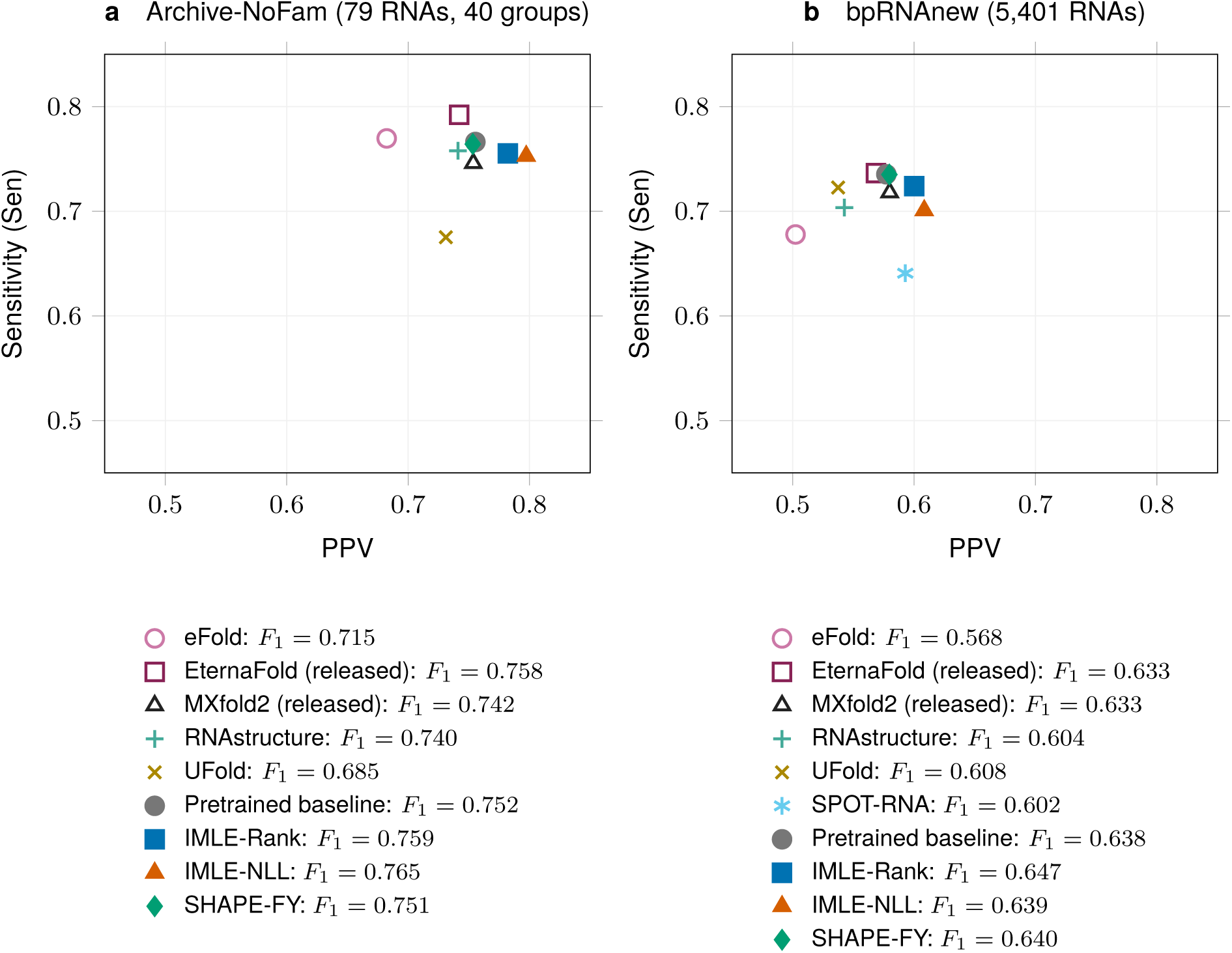
Precision and sensitivity on external structural tests. Points show mean precision (PPV) and sensitivity (Sen); legend values give directly averaged base-pair *F*_1_. **a**, Archive-NoFam: 79 RNAs in 40 structural groups, with equal weight per group. **b**, bpRNAnew: 5,401 RNAs, with equal weight per RNA. Both panels use identical axis limits. The four models trained here use the same five-epoch pretrained baseline, with three Ribonanza and three pri-miRNA/human mRNA epochs for chemical training. Released predictors retain their own training and decoding. SPOT-RNA is shown only in panel b because its available Archive-NoFam results use a different test population (Supplementary Note 5.3).

The bpRNAnew benchmark distributed with MXfold2^1^ reveals a precision–sensitivity trade-off after chemical training (Figure 4b). IMLE-Rank increased mean per-RNA *F*_1_ from 0.638 to 0.647, with precision rising from 0.577 to 0.600 and recall falling from 0.735 to 0.724. IMLE-NLL reached precision 0.608, recall 0.701, and *F*_1_ = 0.639. SHAPE-FY reached 0.640. Precision is defined for 5,401, 5,396, 5,371, and 5,401 RNAs for the pretrained model, IMLE-Rank, IMLE-NLL, and SHAPE-FY, respectively. Every RNA contributes to recall and *F*_1_ (Supplementary Table 10).

## Discussion

Chemical weak supervision can improve complete base-pair prediction while preserving discrete folding constraints and sequence-only inference. With structural pretraining and replay, IMLE-Rank increased TestSetB *F*_1_ from 0.609 to 0.649 in the 2A3 simulations. Following bpRNA–RNAcentral pretraining, all three objectives improved *F*_1_ on ArchiveII and viral RNAs (Figure 3b,c). These examples show that incomplete pairing-state observations can contribute to learning the partners needed for structure prediction.

The three objectives provide alternative ways to use those incomplete observations. I-MLE supplies a surrogate score gradient from discrete predictions and their feature counts^18^. This permits a distribution-free ranking objective or the Wu-model likelihood to supervise the same folding algorithm ^12^. SHAPE-FY trains on a complete assisted-folding target using a Fenchel–Young score comparison ^9,17^. Their different outcomes show that the choice of observation loss matters even with a shared predictor.

IMLE-Rank provides a practical option for adapting pretrained models to heterogeneous probing data. It improves TestSetB accuracy with either synthetic probe and after RNAgym training, and gives the largest bpRNAnew improvement among the three objectives. Its loss is unchanged by a strictly increasing transformation of the retained reactivities, reducing dependence on absolute scales that vary with assay processing and experimental conditions^5,7,8^. IMLE-Rank performs similarly with the two synthetic probes despite their different observation coverage, whereas IMLE-NLL is more sensitive to the probe. These properties motivate its use across heterogeneous profiles. Controlled changes in measurement noise and missingness would quantify its robustness to data quality.

Chemical probing can expand the training sets of structure predictors to include RNA classes with few complete structure annotations. For example, our fine-tuning uses reactivity profiles for primary microRNAs and regions of human messenger RNAs ^13^. The simulations show that learning from pairing-state information can improve complete base-pair prediction for related RNAs. The smaller TestSetB gain after RNAgym training is consistent with its fewer detected sequence matches to that test, suggesting that structural gains may depend on which RNA families the probing profiles cover.

The higher AUROC on held-out RNAgym profiles shows that training improves the separation of reactivities between predicted paired and unpaired positions (Table 2). This provides evidence that the model learns sequence-dependent distinctions consistent with experimental pairing-state information. The viral collection has both probing data and structural references for the same RNAs. On this collection, IMLE-Rank improves chemical agreement and exact-pair accuracy (Figure 3c and Supplementary Table 8).

Structural replay may help retain the partner information learned during pretraining. Rehearsing earlier examples can reduce catastrophic forgetting during sequential learning^31,32^. Here, structure- labeled examples continue to reward complete pairing patterns while chemical examples constrain local states. In the simulations, IMLE-NLL gained more from replay with DMS than with 2A3 (Figure 2). DMS also supplied fewer observed positions (Supplementary Table 3). With synthetic 2A3, pretrained adaptation outperformed joint training for both I-MLE objectives, while SHAPE-FY slightly improved with joint training. Replay added optimizer updates, and the joint and adaptation protocols also differed in prior structural exposure and observation filters.

The external benchmarks show how the choice of test RNAs and the balance between precision and sensitivity shape the observed gains. On bpRNAnew, both I-MLE objectives increase precision and reduce sensitivity, with the larger net *F*_1_ gain for IMLE-Rank (Figure 4b). On Archive-NoFam, their group-mean gains on the primary 79 RNAs reverse when all 89 RNAs are included (Figure 4a and Supplementary Table 11). The primary 79 RNAs and bpRNAnew have no full-length exact matches in our training inputs. The separate analysis of the PDB and ArchiveII tests shows that high local identity can occur even without a full-length match (Supplementary Tables 12 and 13).

Discrete inference supplies structural constraints that local reactivities alone cannot provide. The current decoder restricts predictions to pseudoknot-free structures, while thermodynamic scores and structural examples help choose partners. Our objectives connect indirect experimental measurements to this constrained neural predictor. Because prediction uses sequence alone, a probing dataset can inform predictions beyond the RNAs measured in that experiment.

## Methods

### Model and changes to MXfold2

Our extension changes the training objective while retaining the MXfold2 neural network and structure prediction algorithm. The original method uses a structured hinge loss: training favors the reference structure over alternatives by a margin ^1^. We replace this loss with a Fenchel–Young (FY) loss for examples with known structures. For examples with chemical measurements, we either construct a target structure by assisted folding (SHAPE-FY) or apply a reactivity loss to the predicted pairing states through implicit maximum-likelihood estimation (I-MLE).

MXfold2 predicts the structure with the highest combined neural and thermodynamic score. Let *x* = *x*_1_ · · · *x_L_* be an RNA sequence of length *L*, and let S(*x*) be its allowed pseudoknot-free structures: base-pair arcs do not cross when drawn above the sequence. The neural network, with parameters *v*, produces local folding scores *θ* = *h_v_*(*x*). For a structure *y* ∈ S(*x*), each component of *ϕ*(*x, y*) counts how often a local score is used, for example in a stacking or loop term. The scalar *f_T_* (*x, y*) is the negative Turner free energy. We use

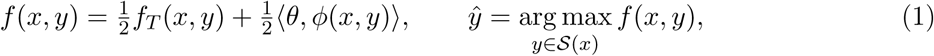

where the inner product sums each local score multiplied by its feature count. The original MXfold2 adds the neural and thermodynamic scores; here we use their equal-weight average. Multiplying the unperturbed sum by a positive constant leaves its maximizing structure unchanged, but the scale must be retained in the losses and pseudoenergy calculations. Turner parameters remain fixed in the primary comparisons. Allowed pairs are AU, CG, and GU in either orientation. All maximizations over structures below are restricted to S(*x*).

Chemical losses use whether each nucleotide is paired, without specifying its partner. Write bp*_i_*(*y*) = 1 if nucleotide *i* is paired in *y*, and 0 otherwise. The decoder returns binary pairing states. To obtain a chemical-loss derivative, we define the loss on a continuous vector *p* ∈ [0, 1]*^L^* and evaluate that derivative at *p* = bp(*y*). The measured reactivity at position *i* is *d_i_*; *d* denotes the full profile. Because different structures can share the same paired positions, agreement with these states does not establish correct base-pair partners.

### Gaussian perturbations of folding scores

Gaussian noise lets training compare alternative structures near the model’s current prediction. For each scalar entry of the raw neural output, we independently draw a standard-normal value *z* ∼ N(0, 1) and add *ɛz*. The model then expands these outputs into the local scores used by folding.

Let *Z* denote the noise after this expansion and let *ɛ* be its fixed scale. The perturbed scores are

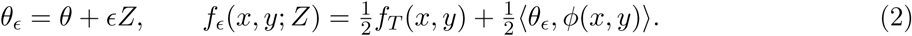

The noise changes the neural scores supplied to the folding algorithm. The network weights and Turner parameters are held fixed during each folding calculation. Supplementary Note 1 gives this construction. We use one noise realization per sequence and keep it unchanged across the I-MLE folding calculations described below. Structural pretraining uses *ɛ* = 0.75.

### Fenchel–Young loss for known target structures

The FY loss trains the model to score a target structure above a competing prediction^17^. Let *y* be a fixed target, and let *y_ɛ_* = arg max*_y_ f_ɛ_*(*x, y*; *Z*) be the perturbed prediction. The loss used for an update and its expected objective are

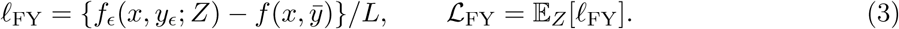

The target is scored without noise, using the same neural-network evaluation. Division by *L* normalizes by sequence length. The expectation is over the noise draws. At each stochastic optimization step, we sample the noise once and use that sample’s loss, rather than calculate the expectation explicitly. For a fixed target, the omitted FY terms do not depend on model parameters and do not affect training. Unlike the original hinge loss, prediction uses Gaussian noise rather than a structure-dependent margin.

The FY gradient is computed from the features of the two structures, without differentiating their pairing decisions. For fixed *y*, the score derivative is *C*(*x, y*) = *ϕ*(*x, y*)*/*2, the feature-count vector weighted by the neural contribution in Equation 1. The score derivative used for backpropagation is therefore

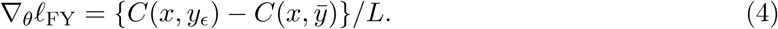

The folding algorithm returns the selected structures and their feature counts. We differentiate the scores of the two selected structures while holding their base-pair choices fixed; we do not differentiate the structure-selection algorithm. Ordinary backpropagation carries this derivative through *θ* = *h_v_*(*x*) to the network parameters *v*. The I-MLE update below uses the same weighted counts *C*(*x, y*). Minimizing the loss raises target features relative to competing features. We use this loss for structural pretraining, structural replay, and the assisted-folding targets described next.

### SHAPE-FY: learning from assisted-folding targets

SHAPE-FY uses reactivities to construct a complete target structure for the FY loss. Assisted folding adds a reactivity-dependent energy correction, or pseudoenergy, to the folding calculation. We use the logarithmic form introduced for SHAPE-directed folding^9^, with slope *m* and intercept *b*, applied as a penalty on each paired position:

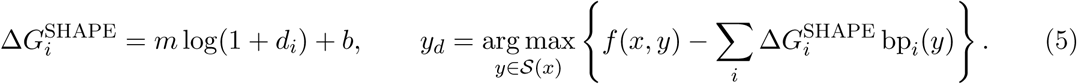

We use *m* = 2.6 and *b* = −0.8. The correction is computed for retained observations with *d_i_ >* −1 and is zero elsewhere. High reactivity discourages pairing; the negative intercept can encourage pairing at low reactivity.

The assisted structure *y_d_* serves as a target, or teacher, for the same MXfold2 model. We remove the pseudoenergy and rescore it without noise, then apply

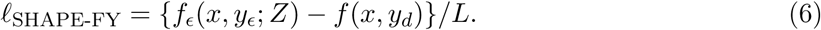

Both scores use the same neural-network evaluation and the weights in Equation 1. We hold *y_d_* fixed during the FY gradient calculation and regenerate it as training proceeds. Reactivities therefore guide training targets, while prediction after training requires sequence alone.

### I-MLE: backpropagating a chemical loss through folding

I-MLE supplies a surrogate gradient for a loss calculated from a discrete predicted structure^18^. A small score change usually leaves the selected structure unchanged, so ordinary differentiation through structure selection gives zero almost everywhere. I-MLE instead repeats folding after shifting the scores in directions determined by the chemical loss. The resulting change in structural features supplies the score gradient passed to the neural network. The chemical loss can be any differentiable function of the continuous pairing-state vector; it need not be a likelihood. The two functions used here are defined in the following subsections.

#### 1. Differentiate the loss with respect to pairing states

We first predict *y*_0_ = arg max*_y_ f_ɛ_*(*x, y*; *Z*). For a chemical loss *ℓ*(*d, p*), we temporarily treat *p* as continuous and calculate

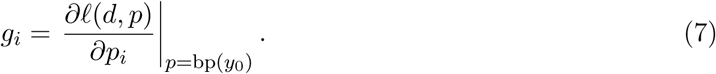

A positive *g_i_* means that lowering the paired indicator at position *i* would decrease the continuous loss locally. Unobserved positions have *g_i_* = 0. This interface accepts a differentiable loss on pairing states; it does not supply a general loss on partner identities.

#### 2. Predict two structures with shifted pairing scores

We convert *g* into positional score shifts Δ*G_i_* = *νg_i_/s*, where *ν >* 0 sets their size. Root mean square (RMS) normalization uses 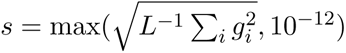, including zeros at unobserved positions. The main comparisons use central differences: after the initial fold *y*_0_ used to calculate *g*, we compute two further folds with opposite score shifts. Both reuse the same Gaussian draw *Z*:

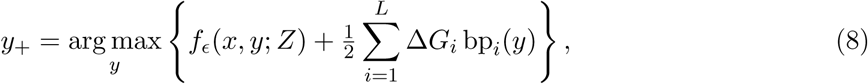

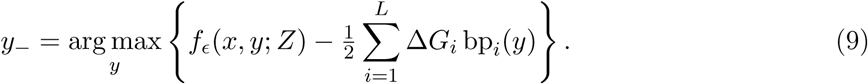

Thus both *y*_+_ and *y_−_* include the loss-directed score shift; the initial *y*_0_ does not. For *g_i_ >* 0, the second fold discourages pairing at *i*, moving toward a lower chemical loss. The first fold shifts the score in the opposite direction. Increasing *ν* increases both shifts and can change which structures are selected.

#### 3. Backpropagate the feature-count difference

The central I-MLE estimate uses the difference between the two structures’ feature counts as a score gradient:

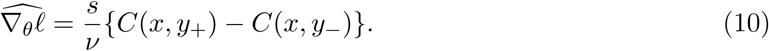

We supply the vector in Equation 10 to neural backpropagation as the chemical-loss gradient with respect to the local scores *θ*. The usual chain rule then gives 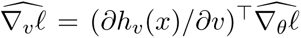 for the network parameters. The loss derivative *g* and its normalization *s* are calculated for the current example; they are not tunable parameters. Together with *Z* and the selected feature counts, they are held fixed during this backward pass. The scale *ν* affects both the selected structures and the factor *s/ν*; it does more than rescale a fixed gradient. If the two folds have identical feature counts, this chemical gradient is zero.

The count difference is a surrogate gradient, not the ordinary derivative of the discrete prediction.

Supplementary Note 1 details the perturbation construction and numerical implementation.

### IMLE-Rank: learning the order of reactivities

IMLE-Rank favors lower reactivity at predicted paired positions than at predicted unpaired positions. It uses a Mann–Whitney objective based on ranks, without fitting a reactivity distribution. Let *M* be the observed positions retained for the loss and *n* = |*M* |. For *i* ∈ *M*, let *R_i_* be the ascending rank of *d_i_*, starting at 1 and averaging ranks for ties. Define *c_i_* = (*n* + 1)*/*2 − *R_i_*, *N_P_* = *_i∈M_ p_i_*, and *N_U_* = ∑*n* − *N_P_* . To calculate the derivative required by I-MLE, we first extend the loss to continuous *p_i_* ∈ [0, 1]. For *n* ≥ 2, the extension is

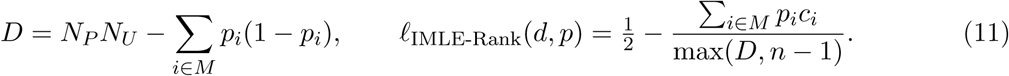

At a binary prediction containing both paired and unpaired observed positions, the loss equals one minus AUROC, defined using unpaired states as positive labels and reactivities as scores. Lower reactivity in the paired group counts as agreement, with half credit for ties. The term *∑_i_ p_i_*(1 − *p_i_*) is zero at a binary prediction, but its derivative with respect to *p_i_* is 1 − 2*p_i_*, which is nonzero at 0 and 1. It can therefore change the I-MLE gradient and is retained when differentiating the loss. This continuous denominator excludes comparisons of a position with itself. Its floor gives a training loss of 0.5 and finite derivatives when all observed positions have the same pairing state. The evaluation AUROC is undefined in this case; the training convention does not fill that missing evaluation value. Constant reactivities give zero gradient; fewer than two observations give zero loss and gradient. The derivative of this loss supplies *g* in Equation 7.

### IMLE-NLL: learning with the Wu reactivity model

IMLE-NLL models the reactivity distribution separately for paired and unpaired nucleotides. Following the SHAPE observation model of Wu et al. ^12^, we use a generalized extreme-value (GEV) density *p_P_* (*d*; *u*) for paired positions and a Gamma density *p_U_* (*d*; *u*) for unpaired positions. The vector *u* contains their learned parameters. For a nonempty set of retained positions *M*, the negative log likelihood is

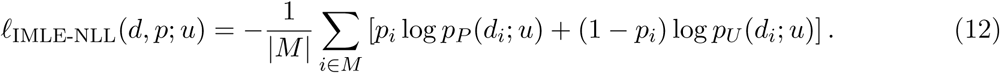

At a binary prediction, each position contributes the log density for its predicted pairing state. The derivative with respect to *p_i_* is [log *p_U_* (*d_i_*; *u*) − log *p_P_* (*d_i_*; *u*)]*/*|*M* |. I-MLE uses this derivative to update the folding scores. The distribution parameters *u* receive their gradients directly by differentiating the likelihood loss, without an I-MLE estimate. We learn separate distributions for each probe or dataset/probe combination. Reactivities are clipped to [0.01, 1], densities have a 10*^−^*^5^ floor, and parameters are bounded. Supplementary Note 1 gives the densities, initial values, and numerical treatment.

IMLE-NLL and EternaFold both relate chemical measurements to pairing states, but use different likelihoods ^14,15^. CONTRAfold-SE and EternaFold use Gamma observation distributions and sum over possible structures when computing the chemical likelihood. IMLE-NLL instead evaluates the Wu-model loss at one predicted structure and uses I-MLE to update the neural scores. Thus our objective is not a likelihood marginalized over structures.

### Retaining structural information during training

Structural replay retains direct structure supervision while the model learns from chemical profiles. Chemical examples use one of the three chemical objectives; replay examples use the FY loss with a complete structural target. The replay fraction is the proportion of all training examples drawn from the structural replay collection. We hold chemical exposure fixed, so higher replay fractions add both structural examples and optimizer updates. RNAgym fine-tuning and fine-tuning on the eFold data traverse shuffled cycles of the structural collection and weight chemical and structural losses by 0.5 and 1, respectively.

Thermodynamic regularization keeps the learned folding scores close to the fixed Turner score. The precise score used in this penalty depends on the objective; Supplementary Note 1 gives the formulas. Table 1 summarizes the optimizer and loss parameters.

### Training data and comparisons

TrainSetA provides 3,166 RNAs for MXfold2 pretraining, structural replay, joint training, and EternaFold training. The main adaptation comparisons use two training settings with different starting models. The first pretrains on TrainSetA for 100 epochs, then learns from simulated Train- SetB profiles or RNAgym measurements. The second pretrains on the larger bpRNA–RNAcentral collection for five epochs, then learns from Ribonanza and mammalian chemical profiles. Within each MXfold2 setting, comparisons among the three chemical objectives use a shared starting model and the same training budget at a given replay fraction. Differences between these settings also involve pretraining data and chemical exposure, so they are not single-factor comparisons. An epoch visits each primary training sequence once; replay adds structure-supervised examples to that exposure.

### Structure collections and pretraining

We use the TORNADO sequence–structure benchmarks introduced by Rivas et al. ^16^ and adopted in the MXfold2 study ^1^. TrainSetA compiles structure- prediction benchmarks from the literature. TrainSetB and TestSetB derive from seed alignments of 22 RNA families in Rfam v10.0^33^, with sequence-level reference structures obtained from alignment consensus annotations. The collections contain 3,166, 1,094, and 430 RNAs, respectively; lengths and total nucleotide counts are given in Supplementary Table 2. The TrainSetA baseline was pretrained for 100 epochs with the FY loss and the AdaBelief optimizer ^34^. Fine-tuning restores the model and its optimizer state, including the moments summarizing previous gradients and the final pretraining update count. Learning rate and weight decay change from 0.001 and 0.01 to 10*^−^*^4^ each; Turner parameters remain fixed. New IMLE-NLL distribution parameters start with zero optimizer moments.

### Target-structure controls

We compare different sources of complete training structures. The reference-structure control uses annotated structures as FY targets. Two other controls use targets from RNAfold-assisted or MXfold2-assisted folding with a teacher that is not updated during student training. SHAPE-FY instead constructs its teacher from the current student scores.

### Synthetic reactivities

We generate synthetic profiles from the annotated TrainSetB pairing states to test learning under a controlled chemical signal. Paired and unpaired values are sampled from GEV and Gamma distributions, then mapped by quantile to empirical distributions for each probe and nucleotide type. These empirical distributions use only RNAgym development training measurements. Observed positions are sampled independently of pairing state within each nucleotide type. The generator produces one profile for the SHAPE reagent 2A3 and one DMS profile per RNA, covering 67,412 and 34,237 positions, respectively. These are structure-derived simulations, not measured weak supervision. Supplementary Table 3 gives the saved observation counts and denominator definitions used for calibration.

Synthetic adaptation evaluates the three chemical objectives and two fixed-teacher FY controls at five replay rates. Each of the five methods uses either 2A3 or DMS profiles, giving ten method–probe combinations; direct supervision with reference structures supplies one further condition. Crossing these eleven conditions with replay fractions of 0, 5, 10, 20, and 50% gives 55 runs of 100 additional epochs after pretraining (update counts in Supplementary Table 4).

### Joint structural and chemical training

MXfold2 joint training uses a common random neural initialization and fresh AdaBelief states. Each joint condition combines the 3,166 structure-labeled TrainSetA RNAs with 1,094 TrainSetB RNAs carrying synthetic reactivities or target structures. The epoch objective is mean structural FY loss plus 0.5 times mean auxiliary loss, each with thermodynamic regularization. Only finite, probe-compatible reactivities greater than 10*^−^*^5^ are retained. The joint conditions and their structural-only control use a fixed 100-epoch endpoint from random initialization. Supplementary Note 2.1 gives the observation filters, controls, normalization, and update budgets.

### RNAgym profiles

RNAgym supplies chemical measurements for training^6^. Its official training split uses 40%-identity sequence clusters and contains 117,773 sequences and 198,411 profiles. Our development training split contains 112,815 sequences and 189,858 profiles; 4,958 sequences are held out by cluster. The measurements use the SHAPE reagents 2A3, 1M7, and NMIA, together with DMS and the chemical-modification reagent CMCT. Probe-specific observation masks are given in Supplementary Note 1. A chemical epoch visits every training sequence and all its profiles. For IMLE-Rank, the retained bases are evaluated separately for A and C under DMS, and together under SHAPE. We average these losses within a profile, then across replicates, experimental conditions, and probes. I-MLE is applied once to the resulting sequence-level loss, using one initial prediction and two shifted predictions. Observation masks and loss aggregation are specified in Supplementary Note 1.

The RNAgym comparison uses the default 20% structural replay and evaluates all three models at the fixed third chemical epoch against their shared baseline. IMLE-Rank and IMLE-NLL use central I-MLE with RMS normalization, *ν* = 0.1, and noise scale *ɛ* = 0.1. Twenty-percent replay adds 28,204 structural examples per epoch and gives 35,255 updates. All models use the same pretrained starting point, and the epoch is not selected by test performance.

### EternaFold training

We train EternaFold independently on all 3,166 TrainSetA sequences, using structural supervision alone, synthetic TrainSetB 2A3 profiles, or official RNAgym training profiles ^15^. Every model starts from zero parameters and jointly optimizes structural and chemical likelihoods when profiles are available. Prediction uses maximum expected accuracy decoding with pairing-weight parameter *γ* = 6. Supplementary Note 3 specifies the structural preprocessing, observation filters, likelihood weights, and optimizer. The EternaFold models trained here are distinct from the released EternaFold predictions used in the larger structural benchmark.

#### bpRNA–RNAcentral pretraining and chemical fine-tuning

The larger-data experiment uses the MXfold2 extension with training collections released by the eFold study^13,19,20^. Its 255,767 sequences comprise 45,507 bpRNA examples and 210,260 RNAcentral examples after removing 931 exact duplicates and prioritizing bpRNA labels. As in the eFold study, RNAcentral contributes structures predicted from sequence with RNAstructure^21^, selected from the 211,191 records retained in the public eFold training collection. All three objectives start from the same model after five pretraining epochs, restoring its complete AdaBelief state at the final pretraining step. Training annotations are projected onto canonical, noncrossing structures; test references retain their original pairs. Supplementary Note 4.1 gives pretraining details.

Chemical fine-tuning proceeds in two fixed stages. The first visits all 45,729 sequences in the Ribonanza collection released with eFold for three epochs^13,22^. The second visits 1,096 primary microRNA (pri-miRNA) and 1,375 human messenger RNA (mRNA) sequences for three further epochs. Only measured profiles supervise chemical examples; supplied structure labels for those examples are not used. Twenty-percent structural replay draws from the full 255,767-sequence pretraining collection, including its RNAcentral pseudo-labels. IMLE-Rank, IMLE-NLL, and SHAPE-FY share the same starting checkpoint and learning-rate/weight-decay setting of 10*^−^*^4^, chemical weight 0.5, replay weight 1, and thermodynamic coefficient 0.125. IMLE-Rank and IMLE-NLL use central I-MLE with RMS normalization and *ν* = *ɛ* = 0.1.

The second stage continues the model, complete AdaBelief state, random states, and replay position reached by the first stage. IMLE-NLL also restores its six Wu distribution modules and their optimizer states. New Wu parameters start with zero optimizer moments in the first stage. Resetting the second-stage epoch counter preserves these learned states. We compare the models after all six fine-tuning epochs, without selecting a peak from test scores. Supplementary Note 4 gives the stage-specific data exposure, update counts, and restored states.

### Structural test collections

Archive-NoFam and bpRNAnew evaluate the same pretrained model and its three six-epoch adaptations. These tests therefore compare the same four models as the primary eFold collections. Evaluation uses sequence alone, with the original reference coordinates and exact base-pair matching. Supplementary Note 5 gives the population definitions, metric denominators, and comparator settings.

### Benchmarks released with eFold

The primary comparison evaluates seven models on all 3,795 RNAs in the public eFold test collections: Protein Data Bank (PDB) 355^23^, ArchiveII 3,370^24^, viral fragments 40^25–28^, and long noncoding RNA (lncRNA) 30^13,29^. Evaluation permits up to 2,000 nt; the longest sequence is 1,995 nt. The 1,000-nt training limit and the selected model weights remain fixed. We compare our five-epoch bpRNA–RNAcentral baseline and three sequentially fine-tuned models with eFold inference using released weights and with EternaFold and MXfold2 predictions distributed by the eFold study. The released MXfold2 model has a different training history from our pretrained baseline. Shared test data and scoring do not equalize training data or budgets.

The 3,795 test sequences have no full-length exact matches to our bpRNA–RNAcentral pretraining or chemical inputs. Local similarity remains: at least 90% local identity with at least 80% coverage of both sequences is found against bpRNA for 10 PDB and 161 ArchiveII RNAs. The similarity analysis covers 3,359 ArchiveII sequences up to 1,000 nt, rather than all 3,370 structural test records. Supplementary Note 6 and Tables 12 and 13 report the complete comparisons. The viral and lncRNA references include structures inferred with chemical probing and assisted folding^13^.

Archive-NoFam tests prediction on structures organized into 47 groups^30^. The primary 79- RNA/40-group collection excludes seven bpRNA-overlap groups from the full 89-RNA/47-group collection. A separate 44-RNA/22-group collection applies the union of published exclusion notes.

Group assignments follow the author-preprint supplementary material; the group map and exclusions are fixed across models. All five full-length exact training matches in the 89-RNA collection are already excluded from the primary 79; no primary RNA exactly matches the pretraining/replay or chemical inputs. These are exact-match checks, rather than family filtering. Supplementary Note 5.3 retains the original population definitions and released-model comparisons.

bpRNAnew tests all 5,401 RNAs in the official MXfold2 release ^1^. Lengths range from 33 to 489 nt; 13 RNAs have ambiguous bases and remain in the primary evaluation. There are no full-length exact matches to this pretraining/replay collection or the chemical inputs. All four models use the same prediction and scoring functions on these tests as on the four eFold collections. Supplementary Note 5 gives details. Released predictors retain their own training and decoding settings, detailed in Supplementary Note 5.2^13,15,21,35,36^. Neither bpRNAnew nor Archive-NoFam provides chemical observations for the present evaluation.

### Evaluation metrics

#### Base-pair accuracy

We measure structural accuracy by exact agreement between predicted and reference pairs. Positive predictive value (PPV) is precision, and sensitivity (Sen) is recall. Let *TP* count correctly predicted pairs, *FP* predicted pairs absent from the reference, and *FN* reference pairs missing from the prediction. For each RNA,

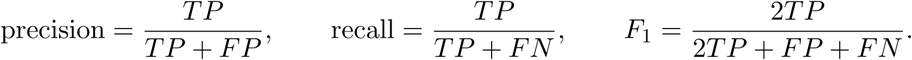

The primary score is the arithmetic mean of per-RNA *F*_1_ values, also called RNA-macro *F*_1_. The overall score for the four eFold collections averages all 3,795 RNA scores equally, giving each collection weight proportional to its size. Dataset-specific treatment of empty predictions and references is detailed in Supplementary Note 5.1. We report their valid RNA counts as *n_P_* and *n_R_*. Every RNA contributes to macro *F*_1_, which is not reconstructed from mean precision and recall. The TestSetB comparisons retain the original benchmark convention: a zero precision or recall denominator gives a score of zero, and all 430 RNAs contribute to each reported mean. The eFold collections and bpRNAnew average only defined precision and recall values. Supplementary Tables 9 and 10 give the plotted means and their valid counts.

Archive-NoFam follows the published benchmark by giving equal weight to each structural group ^30^; this prevents larger groups from dominating the overall result. Each per-RNA metric is first averaged within its structural group, and those group means are averaged with equal weights. This group-macro score gives each structural group the same weight, even when groups contain different numbers of RNAs. We also report the RNA mean separately. For this benchmark, undefined per-RNA precision or recall is assigned zero, following its published convention; this differs from excluding undefined precision in the eFold collections.

### Agreement with reactivity

We evaluate chemical agreement using the unpaired states of a single predicted structure and the reported reactivities. Here, raw reactivity means the value supplied in the released profile, after its original assay processing and before clipping by our training pipeline. MXfold2 supplies the structure maximizing its sequence-based folding score; measured reactivities are used only to evaluate that prediction. Let *p̄_i_* = 1 − bp*_i_*(*y*), so 1 denotes a predicted unpaired position. These predicted states supply the binary labels, and the reactivities supply the scores; reference base pairs do not enter this metric. We retain finite, probe-compatible observations with *d_i_ >* −10. Among these, let *U* and *P* be the predicted unpaired and paired positions. We calculate

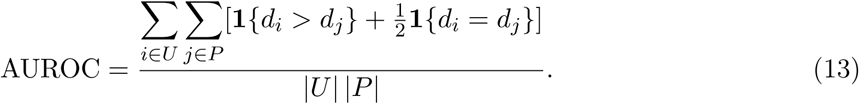

Here **1** is 1 when its condition holds and 0 otherwise. This is the fraction of comparisons in which a predicted unpaired position has higher measured reactivity than a predicted paired position, with half credit for ties. AUROC uses neither base-pair probabilities, reactivity clipping, nor a threshold that divides reactivities into classes.

Spearman’s rank correlation coefficient *ρ* measures agreement between the same binary unpaired states and raw reactivities. We calculate Pearson correlation between the ranks of the unpaired-state vector (*p̄_i_*) and the reactivity vector (*d_i_*), assigning tied values their average rank. Both metrics are undefined when all observed positions have the same predicted pairing state; constant reactivity also makes *ρ* undefined. We exclude these undefined values and do not replace them with a numerical score.

A shared observation mask makes the four-model MXfold2 comparisons use the same measurements for every model. We retain an observation group only when both metrics are defined for all compared models. Chemical evaluation of MXfold2 is restricted to RNAs of at most 1,000 nt. Each DMS profile gives one evaluation on A and one on C. Each SHAPE profile gives one evaluation using all four nucleotide types. We average the retained nucleotide-specific scores within a measurement, then repeated measurements within an RNA, and finally RNAs within a probe. The viral combined score weights all 40 RNAs equally: 25 measured with DMS and 15 with SHAPE. Chemical agreement is evaluated on RNAgym test and viral profiles. Supplementary Note 3.1 and Table 7 give the four-model observation counts.

The MXfold2 RNAgym comparison starts from 433,539 sequences and 702,609 profiles, selecting the highest signal-to-noise ratio (SNR) profile for each RNA and probe. Its four-model intersection at the default 20% replay contains 366,104 2A3 and 318,659 DMS RNA units. For each metric, full-test scores weight each shard mean by its number of RNAs with a defined metric. This reconstructs the RNA mean rather than giving every shard equal weight.

The EternaFold rows in Table 2 use all retained replicate profiles for 2A3 and DMS from the official test measurements, without SNR or sequence-length filtering. Both metrics must be defined in each model, and repeated profiles are averaged within each RNA before the RNA mean is calculated. The structural-only model retains 374,501 2A3 and 328,108 DMS RNAs; the joint model retains 374,303 and 327,784, respectively. Supplementary Table 7 gives profile and nucleotide-group counts. These model-specific sets and replicate processing differ from the common-observation, single-profile MXfold2 comparison, so the chemical columns do not provide a matched six-model comparison. EternaFold decoding uses probabilities internally, while this evaluation uses the paired and unpaired states of its selected structure.

### Training–test sequence similarity

We screened TestSetB against TrainSetB and the official RNAgym training collection with forward MMseqs2 nucleotide searches^37^. We counted an RNA as having a match when an alignment had at least 40% identity and at least 80% coverage of both sequences. For the simulations, we compare mean *F*_1_ changes within the TestSetB groups with and without a match to TrainSetB (Supplementary Note 2).

For the eFold benchmarks, we queried the 355 PDB and 3,359 ArchiveII RNAs of up to 1,000 nt. We compared them with the 45,507 bpRNA training sequences, the full RNAcentral synthetic source, and its subset retained in public eFold training ^13,20^. Supplementary Note 6 gives the collection definitions, including the relation of retained RNAcentral records to our training inputs.

For query *q* and target *t*, full-length similarity is 1 − *d*_Lev_(*q, t*)*/* max(|*q*|, |*t*|), where *d*_Lev_ is Levenshtein edit distance with unit costs and |*q*| and |*t*| are sequence lengths. We report each query’s maximum over all targets in a collection (Supplementary Table 12). For local similarity, we report identity from forward MMseqs2 nucleotide alignments at 80% query coverage or 80% coverage of both sequences. Full-query containment is counted separately (Supplementary Table 13). The eFold study used the Basic Local Alignment Search Tool (BLAST) for its sequence filtering.

### Statistical analysis

Reported scores are descriptive averages from one training run per condition. Adaptations within each protocol share their starting checkpoint. Repeated chemical measurements are averaged within each RNA, the unit of chemical evaluation. Results are displayed to three significant figures, and numerical source data retain their original precision.

### Use of AI tools

OpenAI Codex assisted manuscript drafting and editing, preparation of figure-generation scripts, and consistency checks against existing numerical results. The experimental results reported here come from the completed training and evaluation runs described above.

## Data availability

Study datasets, numerical source data for the figures and tables, and experimental results are provided in the MXfold2 data repository (https://github.com/mxfold/mxfold2-data/). Access to this repository requires permission. The source RNAgym data and datasets released with eFold are available from https://github.com/MarksLab-DasLab/RNAGym and https://github.com/r ouskinlab/efold_data, respectively. Their versions and processing are specified in Methods and Supplementary Notes 3.1 and 4.1.

## Code availability

The MXfold2 extension is available on the master branch of the MXfold2 repository (https://github.com/mxfold/mxfold2). The analysis scripts are provided in the MXfold2 data repository.

## Acknowledgements

This work was supported by JST GteX [JPMJGX23B4 to K.S.], JST CREST [JPMJCR23N1 to K.S.], and JSPS KAKENHI [JP24H00737 and 25H01166 to K.S.].

## Author contributions

K.S. designed the study. R.Y. and K.S. implemented the methods and performed the experiments.

S.T. contributed to the implementation. R.Y. wrote the initial manuscript draft. All authors contributed to writing and reviewing the manuscript.

## Competing interests

The authors declare no competing interests.

## Notes

https://github.com/mxfold/mxfold2/

### Competing Interest Statement

The authors have declared no competing interest.

