## Supplementary Information for "Learning RNA secondary structure from chemical probing with MXfold2"

Rintaro Yamauchi[1], Sho Tsukiyama[1], Kengo Sato[1,2,3*]

[1]School of Life Science and Technology, Institute of Science Tokyo, Japan

[2]Future Intelligence Visionary Initiative, Institute of Science Tokyo, Japan

[3]AI-Science Nexus Center, Institute of New Industry Incubation, Institute of Science Tokyo, Japan

This supplement provides the methods, numerical results, and evaluation details needed for the comparisons in the main text. Scores are on a 0–1 scale and are displayed to three significant figures. Comparisons retain their specified training conditions, test populations, and averaging rules.

### Supplementary Note 1 Implementation and observation models

#### 1.1 Gaussian score perturbations

Gaussian draws are added to the neural output tensors before their assembly into local folding scores. Let $a_v(x)$ collect the paired, unpaired, length, and enabled additional score tensors, and let $\mathcal{A}_x$ be their linear assembly into the score vector. Independent draws $z_k \sim \mathcal{N}(0,1)$ give

$$\theta = \mathcal{A}_x(a_v(x)), \qquad Z = \mathcal{A}_x(z), \qquad \theta_\epsilon = \mathcal{A}_x(a_v(x) + \epsilon z) = \theta + \epsilon Z.$$

Assembly uses indexing, reuse, and sums, so the entries of $Z$ can be correlated and need not have unit variance. Each reported update uses one noise realization per sequence, shared by the initial and shifted implicit maximum-likelihood estimation (I-MLE) folds. The central count difference includes the 0.5 neural-score weight in the mixed score. Selected structures, feature counts, normalization factors, and loss-directed shifts are held fixed during the surrogate backward pass. Network parameters receive the resulting local-score gradient by ordinary backpropagation. The SHAPE-FY correction is applied at individual paired positions and also to DMS profiles. This differs from the original stacking-based SHAPE calibration [1]. The neural architecture and folding grammar otherwise follow MXfold2 [2].

#### 1.2 Wu-model likelihood and numerical treatment

The Wu observation model uses a generalized extreme-value (GEV) density for paired positions and a Gamma density for unpaired positions [3]. For a reactivity $d$, let $u = (\xi, \mu, \sigma, \alpha, \beta)$ denote GEV shape, location, and scale, followed by Gamma shape and rate. With $t = 1 + \xi(d - \mu)/\sigma$,

$$\begin{aligned} p_P(d; u) &= \sigma^{-1} t^{-1-1/\xi} \exp(-t^{-1/\xi}), && t > 0, \\ p_U(d; u) &= \frac{\beta^\alpha}{\Gamma(\alpha)} d^{\alpha-1} \exp(-\beta d), && d > 0. \end{aligned}$$

Here $\Gamma$ is the gamma function. The initial vector is $(0.774, 0.078, 0.083, 1.006, 1.404)$. Training clips reactivities to $[0.01, 1]$, bounds each GEV parameter to $[0.01, 2]$, bounds the Gamma parameters to $[0.01, 5]$, and floors densities at $10^{-5}$. These operations apply to the training likelihood, not the chemical evaluation.

RNAgym uses direct density evaluation. Fine-tuning on the eFold data instead evaluates the GEV density in log space, with $h = -\log(t)/\xi$ and $\log p_P = -\log\sigma + (1+\xi)h - \exp(h)$. It floors the log density at $\log(10^{-5})$, uses the smallest positive representable value in the support logarithm, and returns the floor outside support or when $h > 10$. The observation clipping and parameter bounds remain the same. Distribution parameters receive direct likelihood gradients; I-MLE estimates the gradient to the folding scores. RNAgym has five probe-specific parameter vectors. The sequential chemical protocol creates six dataset/probe modules, four of which receive observations.

#### 1.3 Observation masks and loss aggregation

RNAgym training first retains finite measurements greater than $-900$, then applies the loss-specific filter: values greater than $-10$ for IMLE-Rank and IMLE-NLL, and greater than $-1$ for the logarithmic SHAPE-FY correction. Negative background-corrected observations are retained until these steps. Evaluation uses the distinct raw-reactivity mask defined in the main text.

Probe aggregation respects the nucleotide specificity of each reagent. DMS ranks are computed separately for A and C, CMCT ranks separately for G and U, and SHAPE reagents use all four nucleotide types. Losses average nucleotide groups within a profile, replicate profiles within an experimental condition, conditions within a probe, and probes within a sequence. A single central I-MLE estimate is applied to that sequence-level loss. Constant reactivities or fewer than two observations give zero IMLE-Rank gradient. Neither these training conventions nor the finite-denominator training value of 0.5 fills an undefined evaluation score.

#### 1.4 Thermodynamic regularization

Thermodynamic regularization penalizes differences between a reference score and the fixed Turner score of the same structure. For a reference $y_{\text{ref}}$ and its score $V$, the penalty is

$$R_T = \frac{C_1}{L}\{V - f_T(x, y_{\text{ref}})\}^2, \qquad C_1 = 0.125.$$

For structural FY, $y_{\text{ref}} = \bar{y}$ and $V = f(x, \bar{y})$. For SHAPE-FY, $y_{\text{ref}} = y_d$ and $V = f(x, y_d)$, with the chemical correction removed. For central I-MLE, $y_{\text{ref}} = y_-$ and $V$ is the full shifted score used to select $y_-$ described in Methods, including Gaussian noise and the negative half-shift. Thus the regularized score differs across losses. The chemical loss weight multiplies both the chemical objective and this penalty. Optimizer weight decay is a separate regularizer.

#### 1.5 Shared optimization settings

Supplementary Table 1: Optimizer and loss settings for the reported MXfold2 comparisons. Pretraining and joint training start with fresh optimizer states; chemical adaptation restores the preceding model and optimizer states. A loss weight applies to its thermodynamic penalty as well. The loss derivative $g$ and its RMS normalization $s$ are computed from each current example, rather than selected as hyperparameters.

| Protocol | Epochs | Learning rate | Weight decay | Accumulation |
|---|---|---|---|---|
| TrainSetA pretraining | 100 | $10^{-3}$ | $10^{-2}$ | 1 |
| bpRNA–RNAcentral pretraining | 5 | $10^{-3}$ | $10^{-2}$ | 1 |
| Synthetic adaptation | 100 | $10^{-4}$ | $10^{-4}$ | 4 |
| RNAgym adaptation | 3 | $10^{-4}$ | $10^{-4}$ | 4 |
| Ribonanza then mammalian adaptation | $3 + 3$ | $10^{-4}$ | $10^{-4}$ | 4 |
| Joint training from random weights | 100 | $10^{-4}$ | $10^{-4}$ | 4 |

| Parameter | Value |
|---|---|
| Neural / Turner score weights | 0.5/0.5 |
| Pretraining/replay FY noise scale | $\epsilon = 0.75$ |
| Chemical/target FY noise and I-MLE shift | $\epsilon = 0.1$, $\nu = 0.1$ |
| Independent noise realizations per sequence | 1 |
| RMS denominator floor | $10^{-12}$ |
| Chemical / structural loss weights | 0.5/1 |
| Thermodynamic coefficient | 0.125 |
| Replay in later adaptation comparisons | 20% |
| SHAPE-FY slope / intercept | $m = 2.6$, $b = -0.8$ |
| IMLE-NLL observation clipping | $[0.01, 1]$ |
| IMLE-NLL density floor | $10^{-5}$ |
| AdaBelief moment coefficients | $(0.9, 0.999)$ |
| AdaBelief numerical stabilizer | $10^{-16}$ |
| Dropout | 0.5 |
| Gradient clipping | Disabled |

Structural pretraining uses a Fenchel–Young (FY) loss, AdaBelief, learning rate 0.001, weight decay 0.01, score perturbation 0.75, dropout 0.5, and thermodynamic coefficient 0.125. Chemical adaptation restores network optimizer moments and step counts, changes learning rate and weight decay to $10^{-4}$, and accumulates up to four examples per update. An incomplete accumulation group is averaged by its actual size. AdaBelief uses moment coefficients $(0.9, 0.999)$, stabilizer $10^{-16}$, and decoupled weight decay[4]. Chemical and replay loss weights are 0.5 and 1. Turner parameters remain fixed; gradient clipping is disabled. New likelihood parameters begin with zero optimizer moments.

The TrainSetA baseline has 100 pretraining epochs and optimizer step 316,600. Synthetic and RNAgym adaptations use seed 0. The primary bpRNA–RNAcentral model uses pretraining seed 1234 and five pretraining epochs; its chemical adaptations use seed 0. I-MLE uses central differences, root mean square (RMS) normalization, and $\epsilon = \nu = 0.1$. Structural replay uses FY perturbation 0.75. SHAPE-FY uses slope 2.6 and intercept $-0.8$. Checkpoints preserve network and likelihood parameters, optimizer state, replay position, and random-number states. Comparisons with the starting checkpoint assess the complete chemical-plus-replay adaptation; a replay-only control matched to that adaptation budget is not available. The primary sequential fine-tuning runs use PyTorch 2.7.1 with CUDA 12.6. Their long short-term memory network uses the native PyTorch backend so that dropout random-number states can be restored, while retaining the MXfold2 architecture and dropout probability.

Supplementary Table 2: Structural collections used for TrainSetB simulations and RNAgym adaptation. TrainSetA supplies pretraining and replay, TrainSetB the structures used to generate synthetic profiles, and TestSetB the structural test.

| Collection | RNAs | Nucleotides | Length range (nt) |
|---|---|---|---|
| TrainSetA | 3,166 | 630,279 | 10–734 |
| TrainSetB | 1,094 | 112,398 | 27–237 |
| TestSetB | 430 | 52,097 | 27–244 |

#### Supplementary Note 2 Synthetic profiles and complete replay comparison

Synthetic profiles are generated from the annotated TrainSetB pairing states with seed 20261020 [5,6]. Observation positions are sampled uniformly within nucleotide type and independently of pairing state. Paired latent values use $\mathrm{GEV}(0.774, 0.078, 0.083)$, with SciPy shape convention $-0.774$; unpaired values use Gamma shape 1.006 and rate 1.404. Within each RNA and nucleotide type, the $n$ sampled latent values are ranked. The zero-based rank $k$ receives quantile $(k + U_k)/n$, where $U_k$ is an independent uniform draw on $[0, 1]$. These quantiles are mapped to empirical probe- and nucleotide-specific distributions fitted only to RNAgym development training data.

Each RNA receives one 2A3 and one DMS profile. There are 67,412 observed 2A3 positions and 34,237 DMS positions; missing or ambiguous positions receive $-999$. The generator uses reference structures and shares distributional assumptions with IMLE-NLL. It neither establishes success with measured weak supervision nor reproduces all experimental missingness patterns.

Supplementary Table 3: Observation coverage used to calibrate the simulations. Counts for RNAgym are from its development training split, with repeated profiles contributing their own nucleotide positions. Retained values are finite, probe-compatible, and greater than $-900$, before the loss-specific filters used during training. The denominator includes all nucleotide positions unless A/C is specified. DMS observes only adenine and cytosine; its percentage over all positions is therefore different from coverage among its chemically eligible positions. Synthetic coverage counts structure-derived profiles, rather than experimental measurements.

| Population | Probe | Denominator | Observed | Positions | Coverage (%) |
|---|---|---|---|---|---|
| RNAgym development train | 2A3 | All positions | 10885637 | 18149875 | 60.0 |
| RNAgym development train | DMS | All positions | 4743594 | 15572928 | 30.5 |
| RNAgym development train | DMS | A/C positions | 4743594 | 8608971 | 55.1 |
| Synthetic TrainSetB | 2A3 | All positions | 67412 | 112398 | 60.0 |
| Synthetic TrainSetB | DMS | All positions | 34237 | 112398 | 30.5 |

All 55 runs start from the same TrainSetA epoch-100 baseline and add 100 training epochs. Five chemical/assisted-target methods use each of two probes, and direct reference-structure supervision supplies one additional condition. Each of these eleven conditions is crossed with five replay fractions. Replay counts the additional TrainSetA examples; the reference-structure control already has annotated primary examples. Table 4 gives exposure and update budgets; more replay also adds updates. The two assisted-target controls use fixed RNAfold- or MXfold2-assisted structures, whereas SHAPE-FY regenerates its target from the current model. Every complete-target control uses FY. TestSetB was used repeatedly during development.

To examine whether the IMLE-Rank gains extend beyond detected TrainSetB matches, we grouped the TestSetB RNAs using the forward MMseqs2 screen described in Methods [7]. There were 125 RNAs with at least one match and 305 without a detected match at 40% identity and 80% coverage of both sequences. We averaged per-RNA $F_1$ in each group for the shared pretrained model and the fixed epoch-100 adaptations with 20% replay. For 2A3 the increases were 0.0677 in the group with matches and 0.0278 in the group without detected matches. For DMS the corresponding increases were 0.0425 and 0.0387. Both test groups draw from the same Rfam families as TrainSetB.

The fixed RNAfold teacher uses ViennaRNA 2.7.0 with `-shapeMethod=Dm2.6b-0.8`, applying the stacking-based correction of Deigan et al. [1]. The fixed MXfold2 teacher applies the paired-position correction defined for SHAPE-FY in Methods. Both teachers use the supplied 2A3 or DMS profiles with slope 2.6 and intercept $-0.8$.

Supplementary Table 4: Per-epoch synthetic adaptation budgets. Replay is the fraction of all examples drawn from TrainSetA. Primary exposure stays at 1,094 examples, with gradients accumulated from up to four examples per update.

| Replay | 0% | 5% | 10% | 20% | 50% |
|---|---|---|---|---|---|
| Replay examples | 0 | 58 | 122 | 274 | 1,094 |
| Optimizer updates | 274 | 288 | 304 | 342 | 547 |

Supplementary Table 5: Complete 55-run synthetic comparison after 100 additional epochs from the shared TrainSetA epoch-100 baseline, seed 0. Precision (PPV), sensitivity (Sen), and $F_1$ are TestSetB means over all 430 RNAs under its original zero-denominator convention. Target-structure controls use FY. Replay percentages count additional TrainSetA examples among all training examples; larger fractions also increase update counts.

| Objective or target | Probe | Replay (%) | PPV | Sen | $F_1$ |
|---|---|---|---|---|---|
| IMLE-Rank | 2A3 | 0 | 0.621 | 0.655 | 0.629 |
| IMLE-Rank | 2A3 | 5 | 0.621 | 0.666 | 0.634 |
| IMLE-Rank | 2A3 | 10 | 0.624 | 0.674 | 0.640 |
| IMLE-Rank | 2A3 | 20 | 0.630 | 0.687 | 0.649 |
| IMLE-Rank | 2A3 | 50 | 0.621 | 0.694 | 0.648 |
| IMLE-Rank | DMS | 0 | 0.617 | 0.662 | 0.630 |
| IMLE-Rank | DMS | 5 | 0.616 | 0.669 | 0.633 |
| IMLE-Rank | DMS | 10 | 0.618 | 0.668 | 0.634 |
| IMLE-Rank | DMS | 20 | 0.633 | 0.685 | 0.649 |
| IMLE-Rank | DMS | 50 | 0.617 | 0.693 | 0.645 |
| IMLE-NLL | 2A3 | 0 | 0.630 | 0.626 | 0.618 |
| IMLE-NLL | 2A3 | 5 | 0.636 | 0.638 | 0.627 |
| IMLE-NLL | 2A3 | 10 | 0.642 | 0.644 | 0.632 |
| IMLE-NLL | 2A3 | 20 | 0.642 | 0.670 | 0.647 |
| IMLE-NLL | 2A3 | 50 | 0.641 | 0.695 | 0.659 |
| IMLE-NLL | DMS | 0 | 0.450 | 0.235 | 0.288 |
| IMLE-NLL | DMS | 5 | 0.551 | 0.363 | 0.410 |
| IMLE-NLL | DMS | 10 | 0.546 | 0.376 | 0.415 |
| IMLE-NLL | DMS | 20 | 0.608 | 0.503 | 0.526 |
| IMLE-NLL | DMS | 50 | 0.624 | 0.683 | 0.644 |
| SHAPE-FY | 2A3 | 0 | 0.538 | 0.639 | 0.578 |
| SHAPE-FY | 2A3 | 5 | 0.562 | 0.657 | 0.599 |
| SHAPE-FY | 2A3 | 10 | 0.571 | 0.661 | 0.606 |
| SHAPE-FY | 2A3 | 20 | 0.579 | 0.672 | 0.615 |
| SHAPE-FY | 2A3 | 50 | 0.590 | 0.677 | 0.623 |
| SHAPE-FY | DMS | 0 | 0.567 | 0.609 | 0.578 |
| SHAPE-FY | DMS | 5 | 0.582 | 0.642 | 0.602 |
| SHAPE-FY | DMS | 10 | 0.596 | 0.657 | 0.616 |
| SHAPE-FY | DMS | 20 | 0.597 | 0.667 | 0.622 |
| SHAPE-FY | DMS | 50 | 0.592 | 0.671 | 0.622 |
| RNAfold-assisted targets | 2A3 | 0 | 0.541 | 0.632 | 0.576 |
| RNAfold-assisted targets | 2A3 | 5 | 0.557 | 0.649 | 0.592 |
| RNAfold-assisted targets | 2A3 | 10 | 0.557 | 0.650 | 0.593 |
| RNAfold-assisted targets | 2A3 | 20 | 0.566 | 0.658 | 0.601 |
| RNAfold-assisted targets | 2A3 | 50 | 0.574 | 0.669 | 0.610 |
| RNAfold-assisted targets | DMS | 0 | 0.537 | 0.630 | 0.573 |
| RNAfold-assisted targets | DMS | 5 | 0.549 | 0.643 | 0.585 |
| RNAfold-assisted targets | DMS | 10 | 0.555 | 0.647 | 0.591 |
| RNAfold-assisted targets | DMS | 20 | 0.560 | 0.654 | 0.596 |
| RNAfold-assisted targets | DMS | 50 | 0.564 | 0.661 | 0.601 |
| MXfold2-assisted targets | 2A3 | 0 | 0.580 | 0.666 | 0.613 |
| MXfold2-assisted targets | 2A3 | 5 | 0.581 | 0.669 | 0.615 |
| MXfold2-assisted targets | 2A3 | 10 | 0.586 | 0.671 | 0.618 |
| MXfold2-assisted targets | 2A3 | 20 | 0.593 | 0.679 | 0.626 |
| MXfold2-assisted targets | 2A3 | 50 | 0.592 | 0.680 | 0.626 |
| MXfold2-assisted targets | DMS | 0 | 0.587 | 0.653 | 0.610 |
| MXfold2-assisted targets | DMS | 5 | 0.590 | 0.660 | 0.615 |
| MXfold2-assisted targets | DMS | 10 | 0.593 | 0.664 | 0.619 |
| MXfold2-assisted targets | DMS | 20 | 0.594 | 0.668 | 0.621 |
| MXfold2-assisted targets | DMS | 50 | 0.588 | 0.671 | 0.619 |
| Reference structures | — | 0 | 0.637 | 0.650 | 0.636 |
| Reference structures | — | 5 | 0.628 | 0.651 | 0.631 |
| Reference structures | — | 10 | 0.626 | 0.658 | 0.633 |
| Reference structures | — | 20 | 0.616 | 0.655 | 0.627 |
| Reference structures | — | 50 | 0.609 | 0.671 | 0.631 |

#### 2.1 Joint structural and chemical training from random initialization

MXfold2 joint training uses the 3,166 TrainSetA sequences and their original structural annotations. The runs start from common seed-0 random neural weights, with fixed Turner parameters and fresh AdaBelief moments. Auxiliary examples are the 1,094 TrainSetB sequences with synthetic 2A3 or DMS profiles, fixed assisted targets, or reference structures. The input audit finds 40 full-length sequences shared between TrainSetA and TrainSetB and no exact match from either collection to TestSetB. The reactivity filter retains finite, probe-compatible values greater than $10^{-5}$, giving 51,856 positions for 2A3 and 29,843 for DMS. Objective-specific processing is then applied to these retained values.

Each joint epoch visits the structural and auxiliary examples in shuffled order. The objective is mean structural FY loss plus 0.5 times mean auxiliary loss, with thermodynamic coefficient 0.125 in both terms. AdaBelief uses learning rate and weight decay $10^{-4}$, with gradients accumulated from four examples per update. The per-example backward weights are the epoch update count divided by 3,166 for structural examples and 0.5 times that count divided by 1,094 for auxiliary examples. Joint conditions use 1,065 updates per epoch, compared with 792 for the structural-only control. Thus the coefficients sum to task weights 1 and 0.5 after division by the update count. The twelve conditions use their fixed epoch-100 endpoints. The separate adaptation series first completes 100 structural pretraining epochs, then adds 100 chemical-training epochs with replay. Prior structural exposure, observation filtering, and update budgets differ between these training procedures.

Structural FY uses Gaussian scale 0.75; chemical objectives use 0.1. IMLE-Rank is the Mann–Whitney objective with central I-MLE, root-mean-square normalization, and $\nu = 0.1$. IMLE-NLL learns per-probe Wu distributions with the same central estimator and retains its $[0.01, 1]$ observation clipping. SHAPE-FY regenerates assisted targets from the current model. The two fixed-teacher controls use RNAfold-assisted or MXfold2-assisted targets; only the latter teacher uses a separate model pretrained on all 3,166 TrainSetA structures. The reference-target control uses the annotated TrainSetB structures. These complete-target controls use FY, and no student starts from the teacher weights.

Supplementary Table 6: MXfold2 joint-training results after 100 epochs from random neural initialization. The twelve conditions share the same seed-0 initial neural weights and use 3,166 TrainSetA RNAs. The eleven auxiliary-supervision conditions also use 1,094 TrainSetB RNAs. PPV, sensitivity (Sen), and $F_1$ are means over 430 TestSetB RNAs under the original benchmark convention. Each model uses its fixed epoch-100 endpoint.

| Auxiliary supervision | Probe | PPV | Sen | $F_1$ |
|---|---|---|---|---|
| Structural supervision only | — | 0.574 | 0.658 | 0.606 |
| IMLE-Rank | 2A3 | 0.578 | 0.669 | 0.613 |
| IMLE-Rank | DMS | 0.584 | 0.653 | 0.609 |
| IMLE-NLL | 2A3 | 0.605 | 0.684 | 0.635 |
| IMLE-NLL | DMS | 0.584 | 0.668 | 0.616 |
| SHAPE-FY | 2A3 | 0.586 | 0.675 | 0.621 |
| SHAPE-FY | DMS | 0.576 | 0.667 | 0.611 |
| RNAfold-assisted targets (FY) | 2A3 | 0.572 | 0.668 | 0.610 |
| RNAfold-assisted targets (FY) | DMS | 0.557 | 0.660 | 0.598 |
| MXfold2-assisted targets (FY) | 2A3 | 0.588 | 0.668 | 0.619 |
| MXfold2-assisted targets (FY) | DMS | 0.582 | 0.663 | 0.612 |
| Reference-structure targets (FY) | — | 0.634 | 0.683 | 0.651 |

### Supplementary Note 3 RNAgym and EternaFold comparisons

#### 3.1 RNAgym adaptation and chemical observations

MXfold2 adaptation uses RNAgym profiles[8]: 112,815 training sequences and 189,858 profiles, with 4,958 sequences held out by the official 40%-identity clusters. Each chemical epoch visits every training sequence and its profiles. Twenty-percent replay adds 28,204 structural examples, giving 35,255 updates per epoch. The comparison uses fixed epoch 3 from the shared TrainSetA baseline, with the optimization settings above.

Evaluation uses predicted unpaired states as labels and supplied raw reactivities as scores, retaining finite probe-compatible values greater than $-10$ without clipping. Chemical evaluation averages eligible nucleotide groups within a measurement, repeated measurements within an RNA, and then RNAs within a probe. Both the area under the receiver operating characteristic curve (AUROC) and Spearman correlation must be defined for every model in the four-model MXfold2 comparison. EternaFold uses the model-specific eligible sets described below. All-paired, all-unpaired, or constant-reactivity groups that make a metric undefined are excluded; they are not replaced by numerical scores. RNAgym shard means are weighted by their eligible RNA counts to reconstruct the RNA mean. Table 7 distinguishes RNA units, repeated profiles, and nucleotide groups.

The MXfold2 block uses the 433,539-sequence, 702,609-profile 2A3/DMS benchmark with the highest signal-to-noise ratio (SNR) profile selected per RNA and probe. The EternaFold block instead retains all available replicate profiles for these two probes without SNR or sequence-length filtering, then averages within each RNA. For each EternaFold model, an observation is retained when both metrics are defined in that model. Its structural-only and joint conditions therefore have different eligible observation sets. Table 7 distinguishes their RNA, profile, and nucleotide-group counts. The EternaFold rows do not include the other probes in the larger source collection.

Supplementary Table 7: Eligible observations for the main RNAgym table. MXfold2 uses one profile per RNA and probe, with both metrics defined in all four models. EternaFold retains all available replicate profiles for 2A3 and DMS and averages them within each RNA. Each EternaFold row uses its own observations with both metrics defined. DMS evaluates A and C separately; 2A3 uses all bases. The two protocols and eligible sets remain distinct.

| Comparison | Probe | RNA units | Profiles | Groups |
|---|---|---|---|---|
| MXfold2, four models | 2A3 | 366104 | 366104 | 366104 |
| MXfold2, four models | DMS | 318659 | 318659 | 610415 |
| EternaFold, structures only | 2A3 | 374501 | 390058 | 390058 |
| EternaFold, structures only | DMS | 328108 | 353514 | 704514 |
| EternaFold, joint | 2A3 | 374303 | 389827 | 389827 |
| EternaFold, joint | DMS | 327784 | 352949 | 703150 |

#### 3.2 EternaFold annotation projection and optimization

The three EternaFold conditions retain all 3,166 TrainSetA sequences. A deterministic dynamic program selects a maximum-cardinality subset of each reference's original canonical pairs under the native noncrossing grammar. Paired positions have at least three intervening nucleotides, and single-child internal or bulge loops contain at most 30 unpaired nucleotides; the latter restriction does not limit hairpin length. Projection changes 453 annotations, retaining 166,055 of 167,320 pairs. It adds no pair and excludes no sequence; none of the projected structures is empty.

The conditions independently start from zero parameters and use structures alone, structures plus 1,094 synthetic TrainSetB 2A3 profiles, or structures plus all 117,773 official RNAgym training sequences and 198,411 profiles. The objective is mean structural negative log likelihood plus 0.5 times hierarchical mean chemical marginal negative log likelihood and $\|w\|^2/(2 \times 3166)$, where $w$ is the vector of fitted parameters[9]. Training retains compatible positive observations greater than $10^{-5}$. Chemical likelihood averages replicates within conditions, conditions within probes, probes within sequences, and sequences equally.

Native limited-memory Broyden–Fletcher–Goldfarb–Shanno optimization stops by its small-step rule after 29, 165, and 27 accepted iterations, respectively, before the 200-iteration cap. Sequence-only prediction uses maximum expected accuracy decoding with $\gamma = 6$. TestSetB retains all original coordinates and ambiguous input bases. Its macro precision and sensitivity assign zero for an empty denominator; all 430 RNAs contribute to each mean. These structural values are in the main tables.

### Supplementary Note 4 Primary bpRNA–RNAcentral training protocol

#### 4.1 Pretraining and shared starting state

The primary collection uses data released with eFold[10]: 45,507 bpRNA annotations and 210,260 RNAcentral structures predicted from sequence with RNAstructure[11][12,13]. The latter are selected from the 211,191 records retained with matching identifiers and sequences in public eFold training, removing 931 exact duplicates and giving bpRNA labels priority. TrainSetA/B is not added, and this condition does not apply 80%-identity clustering. Training annotations are projected onto canonical noncrossing structures; test references retain original pairs.

Pretraining uses FY and the settings in Section 1, one example per optimizer update, and seed 1234. All three chemical adaptations restore the same five-epoch network and AdaBelief state at step 1,278,835. This starting point is the focus of a shared-state comparison, rather than a validation-selected optimum.

#### 4.2 Sequential exposure, replay, and restoration

Stage one visits 45,729 Ribonanza sequences for three epochs[10,14]; stage two visits 1,096 primary microRNA (pri-miRNA) and 1,375 human messenger RNA (mRNA) sequences for three further epochs. Ribonanza supplies 44,916 DMS and 37,920 SHAPE profiles; the mammalian examples supply DMS profiles. Chemical-primary structure labels are not used. Replay samples annotated or pseudo-labelled structures from the complete 255,767-sequence pretraining collection.

Twenty-percent replay adds 11,432 structural examples per stage-one epoch and 618 per stage-two epoch. With accumulation four, these give 14,291 and 773 updates per epoch, respectively, for 45,192 updates over the fixed six epochs. The final accumulation group contains one example and is averaged by one. Replay follows continuing shuffled cycles; stage two continues after the preceding 34,296 replay draws. The final optimizer step is 1,324,027.

At the stage boundary, each objective restores its own network, all AdaBelief groups and moments at step 1,321,708, random-number states, and replay position. IMLE-NLL also restores its six Wu modules and their optimizer states. These likelihood parameters begin with zero moments in stage one. Stage-local epoch counters restart while learned states continue. The chemical weight is 0.5, replay weight 1, and thermodynamic coefficient 0.125. Mixed precision and gradient clipping are disabled. The reported endpoint is all six completed fine-tuning epochs, without selecting a test peak.

#### 4.3 Viral chemical agreement

Supplementary Table 8: Viral chemical agreement before and after the six-epoch fine-tuning comparison. All four models share the same five-epoch pretraining checkpoint and use the same 40 RNA measurements. Evaluating adenine and cytosine separately gives 50 dimethyl sulfate (DMS) evaluations from 25 RNAs; selective 2′-hydroxyl acylation analyzed by primer extension (SHAPE) gives 15 evaluations from 15 RNAs. The area under the receiver operating characteristic curve (AUROC) and Spearman correlation $\rho$ compare predicted unpaired states with raw reactivities. The combined score is the mean over all 40 RNAs. Both metrics are defined for every retained nucleotide group in every model; undefined groups are excluded consistently.

| Model | All RNAs AUROC | All RNAs $\rho$ | DMS AUROC | DMS $\rho$ | SHAPE AUROC | SHAPE $\rho$ |
|---|---|---|---|---|---|---|
| Pretrained baseline | 0.810 | 0.517 | 0.855 | 0.586 | 0.735 | 0.402 |
| IMLE-Rank | 0.836 | 0.563 | 0.876 | 0.624 | 0.768 | 0.461 |
| IMLE-NLL | 0.845 | 0.579 | 0.889 | 0.645 | 0.772 | 0.468 |
| SHAPE-FY | 0.814 | 0.523 | 0.855 | 0.586 | 0.746 | 0.420 |

### Supplementary Note 5 Structural evaluation and external tests

#### 5.1 Metric conventions and figure coordinates

The scatter plots report model-level mean precision (PPV) and sensitivity (Sen). Their legend $F_1$ values average individual $F_1$ scores rather than taking the harmonic mean of the plotted coordinates. Table 9 gives the seven-model comparison on all 3,795 PDB, ArchiveII, viral, and lncRNA references. Precision averages RNAs with at least one predicted pair, and sensitivity averages RNAs with at least one reference pair. Their valid counts are $n_P$ and $n_R$. Every RNA contributes to $F_1$, with a value of one when both pair sets are empty. Original reference coordinates and exact pair matching are retained.

The TestSetB tables use a different, original benchmark convention: undefined precision or sensitivity contributes zero, and all 430 RNAs are averaged for each score. This convention is used consistently for MXfold2 and EternaFold. The RNAgym-trained IMLE-NLL model has one empty TestSetB prediction. These means are distinct from the defined-denominator means on the eFold collections and bpRNAnew.

Supplementary Table 9: Structural coordinates and legend values for the main-text precision–sensitivity plots on PDB, ArchiveII, viral, and lncRNA. PPV and sensitivity are means over defined per-RNA values; $n_P$ and $n_R$ give their valid counts. Every RNA contributes to mean $F_1$. Fine-tuned models use the shared five-epoch starting point and fixed six-epoch endpoint; released predictors retain their own training and decoding.

| Collection | Model | $n$ | $n_P$ | $n_R$ | PPV | Sen | $F_1$ |
|---|---|---|---|---|---|---|---|
| PDB | eFold | 355 | 354 | 346 | 0.888 | 0.928 | 0.894 |
| ArchiveII | eFold | 3370 | 3370 | 3366 | 0.572 | 0.636 | 0.600 |
| Viral | eFold | 40 | 40 | 40 | 0.704 | 0.753 | 0.726 |
| lncRNA | eFold | 30 | 30 | 30 | 0.459 | 0.432 | 0.442 |
| PDB | EternaFold | 355 | 351 | 346 | 0.883 | 0.910 | 0.881 |
| ArchiveII | EternaFold | 3370 | 3370 | 3366 | 0.573 | 0.636 | 0.600 |
| Viral | EternaFold | 40 | 40 | 40 | 0.750 | 0.805 | 0.774 |
| lncRNA | EternaFold | 30 | 30 | 30 | 0.450 | 0.467 | 0.457 |
| PDB | MXfold2 (released) | 355 | 353 | 346 | 0.914 | 0.928 | 0.905 |
| ArchiveII | MXfold2 (released) | 3370 | 3369 | 3366 | 0.728 | 0.760 | 0.740 |
| Viral | MXfold2 (released) | 40 | 40 | 40 | 0.703 | 0.720 | 0.710 |
| lncRNA | MXfold2 (released) | 30 | 30 | 30 | 0.410 | 0.429 | 0.419 |
| PDB | Pretrained baseline | 355 | 349 | 346 | 0.901 | 0.907 | 0.889 |
| ArchiveII | Pretrained baseline | 3370 | 3370 | 3366 | 0.594 | 0.643 | 0.615 |
| Viral | Pretrained baseline | 40 | 40 | 40 | 0.719 | 0.765 | 0.739 |
| lncRNA | Pretrained baseline | 30 | 30 | 30 | 0.425 | 0.467 | 0.444 |
| PDB | IMLE-Rank | 355 | 341 | 346 | 0.909 | 0.884 | 0.871 |
| ArchiveII | IMLE-Rank | 3370 | 3370 | 3366 | 0.618 | 0.641 | 0.626 |
| Viral | IMLE-Rank | 40 | 40 | 40 | 0.783 | 0.776 | 0.778 |
| lncRNA | IMLE-Rank | 30 | 30 | 30 | 0.463 | 0.473 | 0.467 |
| PDB | IMLE-NLL | 355 | 337 | 346 | 0.916 | 0.871 | 0.865 |
| ArchiveII | IMLE-NLL | 3370 | 3364 | 3366 | 0.624 | 0.628 | 0.621 |
| Viral | IMLE-NLL | 40 | 40 | 40 | 0.823 | 0.781 | 0.800 |
| lncRNA | IMLE-NLL | 30 | 30 | 30 | 0.456 | 0.434 | 0.444 |
| PDB | SHAPE-FY | 355 | 350 | 346 | 0.902 | 0.911 | 0.893 |
| ArchiveII | SHAPE-FY | 3370 | 3370 | 3366 | 0.599 | 0.646 | 0.619 |
| Viral | SHAPE-FY | 40 | 40 | 40 | 0.727 | 0.768 | 0.745 |
| lncRNA | SHAPE-FY | 30 | 30 | 30 | 0.429 | 0.470 | 0.448 |

Table 10 gives the Archive-NoFam and bpRNAnew scatter coordinates. The four in-study models are the same five-epoch pretrained model and fixed six-epoch adaptations throughout. Neither external reference structures nor reactivities enter inference. No chemical metric is assigned to these tests, which do not supply probing profiles.

Supplementary Table 10: Structural coordinates for the Archive-NoFam and bpRNAnew comparisons shown in the main figure. Archive-NoFam uses 79 RNAs in 40 structural groups and group-macro averages; its published convention assigns zero to undefined per-RNA precision/recall. bpRNAnew uses 5,401 RNAs and RNA-macro averages, excluding undefined precision/recall and reporting valid counts $n_P, n_R$. $F_1$ is averaged directly, rather than reconstructed from mean PPV and sensitivity.

| Collection | Model | $n$ | $n_P$ | $n_R$ | PPV | Sen | $F_1$ |
|---|---|---|---|---|---|---|---|
| Archive-NoFam | eFold | 79 | 79 | 79 | 0.682 | 0.770 | 0.715 |
| bpRNAnew | eFold | 5401 | 5400 | 5401 | 0.502 | 0.678 | 0.568 |
| Archive-NoFam | EternaFold (released) | 79 | 79 | 79 | 0.742 | 0.792 | 0.758 |
| bpRNAnew | EternaFold (released) | 5401 | 5401 | 5401 | 0.569 | 0.736 | 0.633 |
| Archive-NoFam | MXfold2 (released) | 79 | 79 | 79 | 0.754 | 0.746 | 0.742 |
| bpRNAnew | MXfold2 (released) | 5401 | 5399 | 5401 | 0.580 | 0.718 | 0.633 |
| Archive-NoFam | RNAstructure | 79 | 79 | 79 | 0.741 | 0.758 | 0.740 |
| bpRNAnew | RNAstructure | 5401 | 5401 | 5401 | 0.543 | 0.704 | 0.604 |
| Archive-NoFam | UFold | 79 | 79 | 79 | 0.731 | 0.675 | 0.685 |
| bpRNAnew | UFold | 5401 | 5401 | 5401 | 0.537 | 0.723 | 0.608 |
| bpRNAnew | SPOT-RNA | 5401 | 5398 | 5401 | 0.593 | 0.641 | 0.602 |
| Archive-NoFam | Pretrained baseline | 79 | 79 | 79 | 0.755 | 0.766 | 0.752 |
| Archive-NoFam | IMLE-Rank | 79 | 78 | 79 | 0.782 | 0.755 | 0.759 |
| Archive-NoFam | IMLE-NLL | 79 | 78 | 79 | 0.797 | 0.753 | 0.765 |
| Archive-NoFam | SHAPE-FY | 79 | 79 | 79 | 0.753 | 0.764 | 0.751 |
| bpRNAnew | Pretrained baseline | 5401 | 5401 | 5401 | 0.577 | 0.735 | 0.638 |
| bpRNAnew | IMLE-Rank | 5401 | 5396 | 5401 | 0.600 | 0.724 | 0.647 |
| bpRNAnew | IMLE-NLL | 5401 | 5371 | 5401 | 0.608 | 0.701 | 0.639 |
| bpRNAnew | SHAPE-FY | 5401 | 5401 | 5401 | 0.580 | 0.735 | 0.640 |

#### 5.2 Released predictors and decoding

Comparator predictions retain different training inputs and decoding procedures. On the four eFold collections, eFold uses released weights and published postprocessing, while EternaFold and MXfold2 use predictions distributed with the eFold study[10]. These EternaFold predictions do not evaluate the three models independently trained here. The distributed MXfold2 TrainSetA/B training collection has full-length exact matches to 97 PDB and 777 ArchiveII references. The primary bpRNA–RNAcentral collection has no exact matches to these tests.

On the external tests, eFold uses released weights; MXfold2 uses its released TrainSetA/B model; EternaFold uses version-1 parameters and maximum expected accuracy decoding with $\gamma = 6$; RNAstructure uses Fold minimum-free-energy decoding. UFold uses released `ufold_train.pt` weights, and SPOT-RNA uses its five-model ensemble with native postprocessing[2,9,11,15,16]. These are comparisons on common test populations, not training-matched controls.

Ambiguous bases follow the released implementations. UFold retains its official test procedure's training mode and per-sequence batch-normalization statistics, with unknown-base stacking energies set to zero; known-base features remain unchanged. eFold and MXfold2 use native unknown-base encodings. The same 13 ambiguous bpRNAnew inputs are retained for every predictor.

##### 5.3 Archive-NoFam populations and group means

Archive-NoFam contains 89 RNAs of 21–339 nt in 47 structural groups[17]. The primary 79-RNA/40-group population removes the seven groups marked for bpRNA overlap. The group assignments follow the author-preprint supplementary material; 50 descriptive headings map to 47 groups by combining Cas9, Cas12, and Cas13 guide RNAs into their respective groups. The final publication describes the same population size, but identity of its supplementary files with the preprint attachments has not been established. The stated grouping is fixed across our comparisons.

Group-macro metrics average RNAs within groups and then groups equally. Undefined per-RNA precision or sensitivity is assigned zero under this benchmark's convention. RNA-macro means instead weight each RNA equally. Table 11 retains both primary and full populations because the direction of the adaptation effect changes between them. It does not substitute one population for the other.

Five complete sequences in the full 89 match the pretraining/replay inputs: `1f1t_A`, `4k27_U`, `4pcj_A`, `4r4v_A`, and `5m0h_A`. All five are already absent from the primary 79, and neither chemical stage exactly matches an external test. These checks normalize case and T/U; they do not remove every related RNA family. The separate author-supplied SPOT-RNA result, group-macro $F_1 = 0.702$, uses 44 RNAs in 22 groups after the union of additional published exclusions. It is not a score on the primary 79-RNA population.

Supplementary Table 11: Archive-NoFam population and averaging sensitivity for the primary four models. Group $F_1$ weights structural groups equally after averaging within groups; RNA $F_1$ weights individual RNAs equally. Each adaptation uses the same five-epoch starting point and six-epoch endpoint. The primary 79-RNA and full 89-RNA populations remain distinct.

| Model | RNAs | Groups | Group $F_1$ | RNA $F_1$ |
|---|---|---|---|---|
| Pretrained baseline | 79 | 40 | 0.752 | 0.759 |
| IMLE-Rank | 79 | 40 | 0.759 | 0.765 |
| IMLE-NLL | 79 | 40 | 0.765 | 0.768 |
| SHAPE-FY | 79 | 40 | 0.751 | 0.757 |
| Pretrained baseline | 89 | 47 | 0.764 | 0.765 |
| IMLE-Rank | 89 | 47 | 0.758 | 0.752 |
| IMLE-NLL | 89 | 47 | 0.759 | 0.753 |
| SHAPE-FY | 89 | 47 | 0.770 | 0.767 |

##### 5.4 bpRNAnew, ambiguous bases, and valid counts

bpRNAnew uses all 5,401 RNAs in the official MXfold2 release, with lengths 33–489 nt[2]. Thirteen have ambiguous bases. References retain original pairs, including pseudoknots, and every reference has at least one pair. An empty prediction therefore contributes zero to $F_1$ and sensitivity, while its undefined precision is omitted from the precision mean. Table 10 reports the valid counts for all 5,401 RNAs. Neither the pretraining/replay collection nor the two chemical stages has a full-length exact match to bpRNAnew.

#### Supplementary Note 6 Training–test sequence similarity

The similarity analysis separates the bpRNA and RNAcentral components of training from the larger public RNAcentral source. Targets are 45,507 bpRNA sequences, all 226,729 unique sequences in the public RNAcentral synthetic source, and the 211,191 records retained with matching identifiers and sequences in public eFold training. The primary model uses the retained records after duplicate removal with bpRNA.

Queries are PDB 355 and ArchiveII 3,359 at lengths up to 1,000 nt; the eleven longer ArchiveII structural tests are outside this analysis.

Full-length similarity is the maximum of $1 - d_{\mathrm{Lev}}(q,t)/\max(|q|,|t|)$ over every target $t$, where edit distance assigns unit costs to insertion, deletion, and substitution. It is distinct from local alignment identity. Table 12 gives its distributions and thresholds for each target collection.

Local comparisons use MMseqs2 nucleotide search[7] with the following settings.

```
-search-type 3 -min-seq-id 0.6 -c 0.8 -cov-mode 2 -alignment-mode 3 -max-seqs 50000 -strand 1 -mask
0 -e 1000 -s 7.5
```

Query coverage is at least 80%; the two-sided results also require target coverage of at least 80%. Best hits are selected by identity, then coverage and bit score. Searches retain the public forward orientation. An absent heuristic hit is not assigned zero similarity, and this procedure does not reproduce the original eFold BLAST filtering.

Full-length exact matching requires complete query and target strings to agree; substring containment permits a longer target. The contained PDB queries `2NOR-2D`, `2MNC-2D`, and `8TNS-2D` are 23–24 nt long and give four query–target matches. Table 13 keeps these counts separate from identity/coverage thresholds. At both coverages at least 80% and identity at least 90%, bpRNA has matches to 10 PDB and 161 ArchiveII queries, and retained RNAcentral to 3 and 176. The complete RNAcentral source has 64 ArchiveII full-length exact matches, but none remains in the retained subset. These results neither establish family independence nor, by themselves, identify contamination of the published model.

Supplementary Table 12: Maximum full-length normalized edit similarity to actual bpRNA training (45,507), RNAcentral source (226,729), or RNAcentral retained in public eFold training (211,191). Threshold columns count test RNAs, not alignments. PDB contains 355 RNAs and ArchiveII 3,359 within the length limit.

| Training/source | Test | RNAs | Mean | Median | $\geq .8$ | $\geq .9$ | $\geq .95$ | Exact |
|---|---|---|---|---|---|---|---|---|
| bpRNA | PDB | 355 | 0.648 | 0.629 | 24 | 10 | 2 | 0 |
| bpRNA | ArchiveII | 3359 | 0.664 | 0.659 | 461 | 88 | 3 | 0 |
| RNAcentral source | PDB | 355 | 0.604 | 0.593 | 9 | 6 | 4 | 0 |
| RNAcentral source | ArchiveII | 3359 | 0.742 | 0.730 | 1490 | 1107 | 670 | 64 |
| RNAcentral retained | PDB | 355 | 0.602 | 0.593 | 9 | 0 | 0 | 0 |
| RNAcentral retained | ArchiveII | 3359 | 0.675 | 0.644 | 920 | 42 | 0 | 0 |

Supplementary Table 13: Forward MMseqs2 local sequence matches. Every hit covers at least 80% of the query. The following columns add the stated identity and coverage thresholds; both coverages means query and target coverage. Containment counts queries whose complete sequence occurs inside a target, including full-length exact matches. An absent hit does not imply zero similarity.

| Training/source | Test | Query coverage $\geq 80\%$ All hits | Query coverage $\geq 80\%$ Identity $\geq 90\%$ | Both coverages $\geq 80\%$ Identity $\geq 80\%$ | Both coverages $\geq 80\%$ Identity $\geq 90\%$ | Both coverages $\geq 80\%$ Identity $\geq 95\%$ | Contained |
|---|---|---|---|---|---|---|---|
| bpRNA | PDB | 46 | 31 | 21 | 10 | 4 | 3 |
| bpRNA | ArchiveII | 1070 | 166 | 560 | 161 | 23 | 0 |
| RNAcentral source | PDB | 35 | 30 | 9 | 6 | 6 | 6 |
| RNAcentral source | ArchiveII | 1675 | 1260 | 1547 | 1258 | 940 | 106 |
| RNAcentral retained | PDB | 23 | 15 | 9 | 3 | 0 | 1 |
| RNAcentral retained | ArchiveII | 1447 | 176 | 1149 | 176 | 13 | 0 |
